# TACC3-driven translation reprogramming dictates susceptibility or tolerance to mitotic stress

**DOI:** 10.64898/2026.09.18.752750

**Authors:** Ozge Saatci, Aldo Hernández-Corchado, Adriana Aguilar-Mahecha, Breege Howley, Jennifer Rutherford Bethard, Jean-Sebastien Anoma, Josiane Lafleur, Marguerite Buchanan, Elizabeth G Hill, Lauren E Ball, Mathew Sajish, Philip Howe, Mark Basik, Hamed Najafabadi, Ozgur Sahin

## Abstract

Translation reprogramming is central to cancer cell plasticity under stress. However, the molecular players coordinating translation and epitranscriptomic rewiring to determine adaptive responses to mitotic stress remain elusive. Here, we found that microtubule targeting agent (MTA)-induced CDK1 blocks global translation while it phosphorylates and degrades TACC3, a multifunctional adaptor, releasing eIF4A/eIF4E/eIF4G2 initiation factors. This promotes selective m^7^G cap-dependent translation of mRNAs with ‘MTAup’ motif that are functionally required for apoptosis by disrupting proteostasis, promoting MTA-sensitivity. On the other hand, selectively translated TACC3 interacts with eIF3d/eIF4G1 and the m^6^A writer METTL3, mediating switch to m^6^A methylation and m^7^G cap-independent translation of mRNAs with hnRNPC-motif involved in chromosome segregation, driving MTA tolerance. TACC3 inhibition overcomes MTA resistance via restoring translation reprogramming. These findings demonstrate that TACC3 is a pivotal coordinator of translation/epitranscriptomic reprogramming and a therapeutic target in MTA-refractory cancers.

**Highlights:**

- MTA-induced CDK1 blocks global translation while increasing selective translation
- Cap-dependent translation of mRNAs with MTAup motif leads to MTA-induced apoptosis
- TACC3 fuels m^6^A methylation/switch to cap-independent translation and MTA tolerance
- TACC3 serves as a potential therapeutic target to overcome MTA resistance

## Introduction

Translation reprogramming is the central feature of adaptive plasticity^1^, enabling rapid phenotype switching upon cellular stress by favoring efficient and transcript-specific mRNA translation^2^. Mechanisms of translation rewiring is multifaceted and mostly involves alterations in the rate-limiting initiation step executed by the eIF4F initiation complex, a heterotrimeric protein assembly that serves as the central hub for N7-methylguanosine (m^7^G) cap-dependent translation initiation in eukaryotes^3^. eIF4F complex comprises three subunits: the m^7^G cap-binding eIF4E^4^, the helicase eIF4A^5^, and the scaffolding protein eIF4G^6^. Under cellular stresses such as nutrient deprivation, hypoxia, or oxidative stress, the upstream sensor mTORC1 kinase hyper-phosphorylates 4E-binding protein 1 (4E-BP1)^7^, halting the assembly of the heterotrimeric eIF4F complex and blocking the energy-consuming global translation. Despite suppression of global translation, certain transcripts can be selectively translated in a cap-dependent^8^ or -independent manner^9^ to initially mitigate damage, maintain proteostasis and promote survival, or to execute cell death in case of severe or prolonged stress^10^. However, little is known about the functional contribution of cap-dependent and -independent selective translation to response to mitotic stress.

In recent years, tumor cell plasticity emerged as ‘the great escape’ from the growth inhibitory effects of anti-cancer therapies, driving transformation towards a phenotypic state that can tolerate given therapy. Although transcriptomic profiling has provided some insights into the molecular mechanisms underlying acquisition of a ‘drug tolerant’ cell identity^11^, the dynamics of the translational changes during adaptation to drug-induced stress remain largely unknown. Previous studies have demonstrated that transcript abundance does not always reflect protein levels^12^, highlighting the importance of posttranscriptional regulation. Along these lines, mRNA chemical modifications, such as N6-methyladenosine (m^6^A) have recently emerged as key regulators of dynamic gene regulation at posttranscriptional level. m^6^A is the most abundant and conserved epitranscriptomic modification in eukaryotic RNAs^13^ regulating various aspects of RNA biology, including translation and is orchestrated by m^6^A writers, readers, and erasers^14^. However, regulators of the dynamic crosstalk between m^6^A methylation and selective translation during adaptation to drug-induced stress, contributing to drug tolerance and their therapeutic potential to restore drug sensitivity remain to be elucidated.

In this study, we uncover the molecular mechanisms of translation reprogramming governed by specific initiation factors/mRNA sequences and epitranscriptomic mechanisms upon therapy-induced mitotic stress and identified TACC3, a multifunctional adaptor for distinct multi-protein complexes^15^, as the key orchestrator of translation plasticity. We showed that during mitotic stress induced by microtubule targeting agents (MTAs), a cornerstone chemotherapy agent, global translation is suppressed in a CDK1-dependent manner. On the other hand, CDK1-mediated phosphorylation and degradation of TACC3 promotes selective translation of certain mRNAs bearing unique sequence motifs in a cap-dependent manner, leading to apoptotic cell death upon disruption of proteostasis. In tolerant cells, TACC3 is selectively translated mediating the switch to cap-independent selective translation and m^6^A methylation of pro-survival mRNAs. Furthermore, targeting TACC3 with a first-in-class clinical inhibitor overcomes MTA resistance without major toxicity. In sum, our work reveals the novel molecular mechanisms of translation/epitranscriptomic reprogramming triggered by mitotic stress and a novel molecular target, TACC3, orchestrating phenotype switching to drive MTA tolerance/resistance.

## Results

### Mitotic stress-induced decoupling of translation from transcription and suppression of global translation underlies sensitivity to microtubule-targeting agents

To determine the clinically relevant drivers of drug-induced mitotic stress that are potentially associated with response/resistance to MTAs, we first analyzed a published microarray data of docetaxel-based therapy-treated (a mitotic stress inducer) breast tumors from responder vs. non-responder patients^16^ (**Figure 1A**). Network analysis revealed that the MTA-treated responder tumors express lower levels of mitosis- and chromosome segregation-related genes compared to non-responders (**Figure 1B**), potentially related to failure in mitotic progression, as expected. Interestingly, we also observed lower levels of the translation- and RNA metabolism-related gene sets in responder tumors (**Figure 1A, B, Table S1**) that potentially give rise to lower rates of mRNA translation. To analyze the association between MTA response and mRNA translation at the global level in patients, we utilized a second dataset containing RNA-seq and mass spectrometry data of docetaxel-based therapy-treated responder vs. non-responder breast tumors^17^ where translation rate can be inferred by comparing mRNA and protein counts as previously reported^18^. Intriguingly, we observed lower average protein counts in MTA-treated responder tumors despite similar levels of mRNA counts (**Figure 1C**), indicative of a potential suppression of mRNA translation at the global level. These data suggest that translation reprogramming via suppression of global translation could be a novel clinically relevant predictor of susceptibility to mitotic stress.

**Figure 1.**
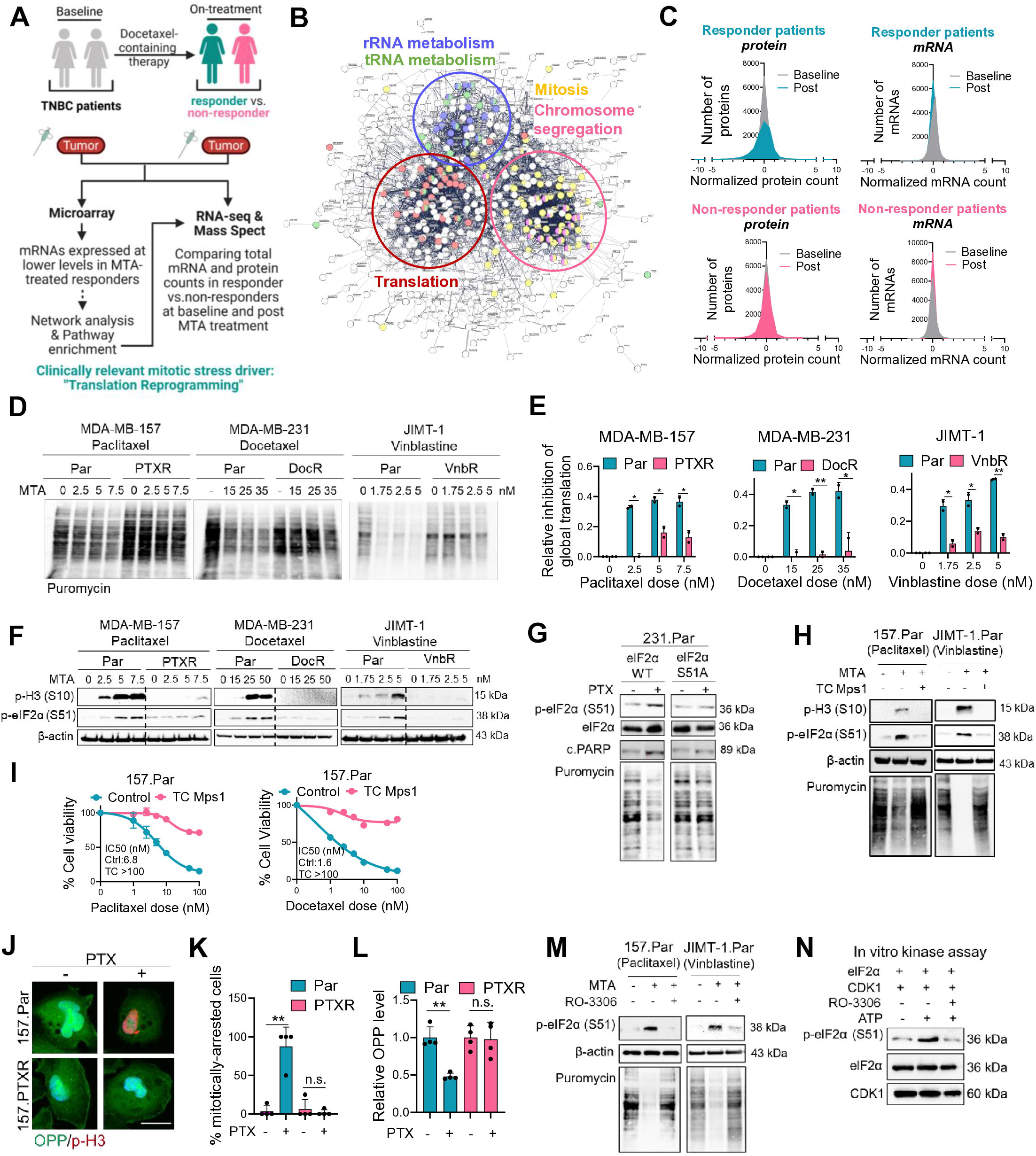
Mitotic stress-induced suppression of global translation underlies sensitivity to microtubule-targeting agents. (A) Pipeline for patient data analysis to identify the clinically relevant drivers of drug-induced mitotic stress. (B) Network analysis among downregulated mRNAs upon MTA treatment in responder vs. non-responder patients. (C) Average protein and mRNA counts in responder vs. non-responder patient tumors before and after MTA-containing therapy. (D) Puromycin labeling in parental (Par) vs. MTA-resistant cells treated with increasing doses of MTAs for 24 hrs. (E) Quantification of puromycin labeling from D (n=2 independent experiments). (F) Western blot (WB) of p-H3 (S10) and p-eIF2α (S51) in cells from D. (G) WB of p-eIF2α (S51) and c. PARP in Par cells overexpressing WT vs. S51A eIF2α and treated with MTAs (50 nM) for 24 hrs. (H) WB of p-H3 (S10) and p-eIF2α (S51), and puromycin labeling in Par cells treated with MTAs (PTX: 10 nM, Vinblastine: 2.5 nM) +/− SAC inhibitor TC Mps1 (5 µM). (I) Percent cell viability in Par cells treated with MTAs +/− TC Mps1 (1 µM) for 72 hrs (n=4-6). (J) OPP and p-H3 (S10) staining in Par vs. PTXR cells treated with PTX (10 nM) for 24 hrs. (K) % mitotically arrested Par vs. PTXR cells from J (n=4). (L) Relative OPP intensity in cells from in J (n=4). (M) WB of p-eIF2α (S51) and puromycin labeling in Par cells treated with MTAs (PTX: 10 nM, Vinblastine: 2.5 nM) +/− CDK1 inhibitor RO-3306 (5 µM). (N) In vitro kinase assay showing CDK1-mediated S51 phosphorylation of eIF2α. PTXR: paclitaxel, DocR: docetaxel, VnbR: vinblastine resistant. Data represent mean ± SD. Two-sided Student’s t-test was used to calculate statistical difference between two groups. \**P*<0.05, ** *P*<0.01; n.s., not significant.

To experimentally determine the effects of MTAs on rates of global protein translation in vitro, we performed puromycin labeling in three different MTA-treated sensitive (i.e., parental) vs. MTA-resistant breast cancer cells (**Figure S1A**) and demonstrated that MTAs suppress global translation only in parental but not in resistant cells (**Figure 1D, E**). Importantly, the phosphorylation of the initiation factor eIF2α that blocks translation initiation^19^ was dose-dependently increased in sensitive cells upon MTA treatment along with the induction of mitotic arrest, which were abrogated in resistance (**Figure 1F**). Mutating the S51 phospho-site restored global translation and reduced MTA-induced cell death (**Figure 1G, S1B**). Since p-eIF2α is also associated with the formation of stress granules^20^, we performed G3BP1 staining (a marker of stress granules) and observed no stress granule formation under MTA doses that induce mitotic arrest, eIF2α phosphorylation and reduction of global translation (**Figure S1C, D**), suggesting that MTA-induced p-eIF2α is likely to be independent of stress granule formation.

Next, we sought to test if global translation blockage is dependent on mitotic arrest, i.e., the hallmark of MTA sensitivity. Induction of mitotic slippage using TC Mps1^21^ reduced MTA-induced p-eIF2α and rescued global translation (**Figure 1H**) as well as cell viability (**Figure 1I**). In situ visualization of nascent protein synthesis using OPP staining demonstrated that it is the mitotically arrested cells that have reduced global translation (**Figure 1J-L**). Since CDK1 is the major mitotic kinase with sustained activity in mitotically arrested cells^22^, we asked if CDK1 could be involved in eIF2α phosphorylation and global translation blockage. Indeed, when CDK1 is inhibited using the CDK1 inhibitor, RO-3306^23^, eIF2α phosphorylation is reduced together with the rescue of global translation (**Figure 1M**). Notably, we showed that CDK1 directly phosphorylates eIF2α in an in vitro kinase assay (**Figure 1N**). Overall, MTA-induced mitotic stress decouples translation from transcription and suppresses global translation via CDK1-dependent eIF2α phosphorylation.

### Selective cap-dependent translation of mRNAs bearing unique sequence motifs disrupts proteostasis and drives mitotic cell death during MTA sensitivity in a CDK1-dependent manner

We found that suppression of global translation upon MTAs was followed by a dose-dependent increase in the phosphorylation of 4E-BP1, a marker of cap-dependent translation^24^, at S83 site that was accompanied by PARP cleavage in sensitive cells but not in resistant counterparts (**Figure S2A**), suggesting that MTAs activate selective cap-dependent translation as a novel mechanism of cell death. These results were further confirmed using the bicistronic reporter system^25^ (**Figure S2A**), and with cap pull down showing increased cap recruitment of initiation factors in MTA-treated sensitive cells (**Figure 2B**). Inhibiting cap complex formation using 4E1RCat or 4EGI significantly reduced apoptosis (**Figure 2C, D**), demonstrating the functional importance of selective cap-dependent translation for MTA-induced cell death. Inhibiting CDK1 (using RO-3306) or inducing mitotic slippage (using TC Mps1) reduced p-4E-BP1 levels and apoptosis and rescued cell viability (**Figure 2D-H, S2B, C**). Notably, inhibiting the canonical 4E-BP1 kinase, mTORC1^26^ using RAD001 or siMTORC1 had no effect on MTA-induced 4E-BP1 phosphorylation (**Figure 2E, G**) and inhibition of cell viability (**Figure 2H, S2D**). We showed direct phosphorylation of 4E-BP1 by CDK1 with an in vitro kinase assay (**Figure 2I**) and demonstrated that mutating S83 phospho-site on 4E-BP1 resulted in reduction of MTA-induced apoptosis (**Figure S2E**). These data demonstrate the functional importance of CDK1-regulated selective cap-dependent translation for MTA sensitivity.

**Figure 2.**
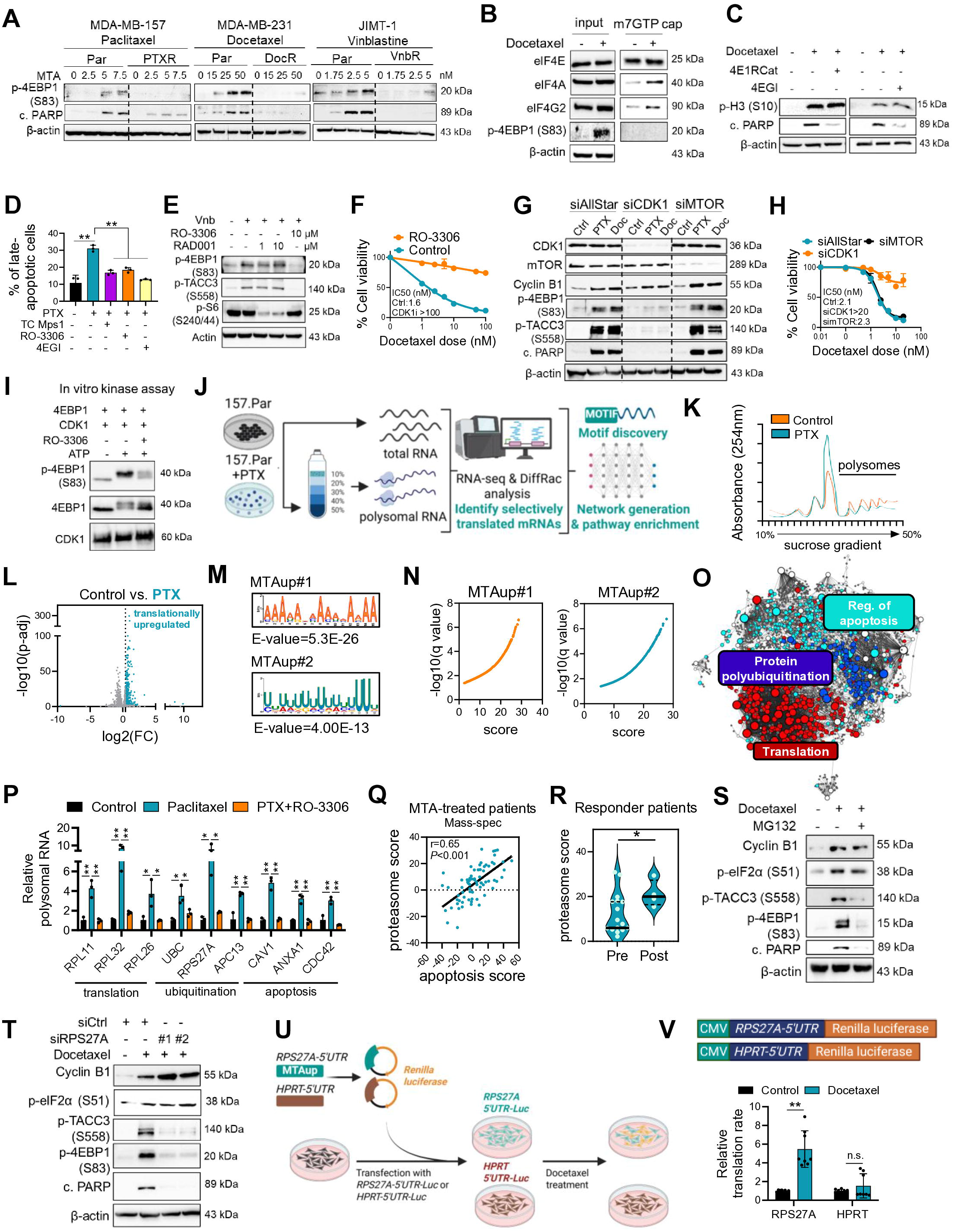
Selective cap-dependent translation of mRNAs bearing unique sequence motifs disrupts proteostasis and drives mitotic cell death during MTA sensitivity in a CDK1-dependent manner. (A) WB of p-4E-BP1 (S83) and c. PARP in Par vs. MTA-resistant cells treated with increasing doses of MTAs for 24 hrs. (B) Cap pull-down in MTA-treated Par cells. (C) WB of p-H3 (S10) and c. PARP in JIMT-1.Par treated with Docetaxel (5 nM) +/− 4E1RCat or 4EGI (50 µM). (D) Percent late-apoptotic cells upon treatment with PTX (50 nM) +/− TC Mps1, RO-3306 (5 µM), or the cap complex inhibitor 4EGI (50 µM) (n=3). (E) WB of p-4E-BP1 (S83), p-TACC3 (S558) and p-S6 (S240/44) in JIMT-1.Par cells treated with vinblastine (2.5 nM) +/− CDK1 inhibitor, RO-3306 (10 µM) or mTORC1 inhibitor, RAD001 (1 or 10 µM). (F) Percent cell viability in 157.Par cells treated with MTAs +/− RO-3306 (n=4-6). (G) WB of p-4E-BP1 (S83), p-TACC3 (S558), Cyclin B1, CDK1, mTORC1 and c.PARP in JIMT-1.Par cells transfected with siCDK1 or siMTORC1 and treated with paclitaxel or docetaxel (5 nM). (H) Percent cell viability in JIMT-1.Par cells transfected with siCDK1 or siMTORC1 and treated with docetaxel (n=3-6). (I) In vitro kinase assay showing CDK1-mediated S83 phosphorylation of 4E-BP1. (J) Scheme of polysome fractionation and RNA-seq from polysome-bound and total RNAs from PTX-treated 157.Par cells. (K) Polysome profiles of 157.Par cells treated with PTX. (L) Volcano plot of differentially translated mRNAs in PTX-treated 157.Par cells. (M) The identified MTAup motifs and their statistical significance. (N) Distribution of MTAup#1 and #2 motif scores across mRNAs selectively translated in sensitive cells. (O) Network analysis among motif MTAup-carrying translationally upregulated mRNAs upon MTA treatment showing enrichment of translation, apoptosis, and protein polyubiquitination. (P) Relative polysomal enrichment of translation, ubiquitination, and apoptosis-related mRNAs in PTX (50 nM)-treated cells with or without RO-3306 (n=3). (Q) Pearson correlation between apoptosis and proteasome scores determined using protein expression in taxane-treated breast cancer patients (n=71). (R) Protein levels of the proteasome score in responder breast cancer patients before and after taxane-based therapy (n=21 pre, n=4 post). (S) WB analysis of mitotic arrest and cell death markers in 157.Par cells treated with Docetaxel (10 nM) with or without 1 µM MG132. (T) WB analysis of mitotic arrest and cell death markers in JIMT-1.Par cells transfected with siRPS27A and treated with Docetaxel (20 nM). (U) Schematic overview of the luciferase reporter assay to determine functionality of MTAup motif for selective translation. (V) Luciferase reporter assay in 231.Par cells transfected with reporter vectors bearing 5’UTR of *RPS27A* or *HPRT* upon 20 hrs of docetaxel treatment (100 nM) (n=8). The relative translation rate was calculated by taking the ratio of the luciferase signal to mRNA levels of the vectors determined by qRT-PCR. In scatter dot plots, the box depicts median, 25th to 75th percentiles, and the whisker depicts min to max for this figure and all others. Data represent mean ± SD. Two-sided Student’s t-test was used to calculate statistical difference between two groups. \**P*<0.05, ** *P*<0.01; n.s., not significant.

To determine the selectively translated mRNAs during MTA sensitivity, we performed polysome fractionation followed by sequencing of polysome-bound vs. total RNAs in MTA-treated sensitive cells (**Figure 2J-L, Table S2**). Polysome fractionation revealed a major reduction of polysome peaks upon MTA treatment (**Figure 2K**), indicative of blockage of global translation^27^. We demonstrated enrichment of translation-related mRNAs among translationally upregulated transcripts in MTA sensitive cells (**Figure S3A**), suggesting a feedforward loop facilitating selective translation. We also demonstrated enrichment of apoptosis- and polyubiquitination-related mRNAs (**Figure S3A**). On the other hand, mitosis and chromosome segregation processes were enriched among translationally downregulated mRNAs, indicative of mitotic failure (**Figure S3B**).

To determine the core network of selectively translated mRNAs, we performed de novo motif discovery analysis. We identified two unique motifs significantly enriched among the 5’UTRs of selectively translated mRNAs in PTX-treated cells (named as MTAup#1 and #2, **Figure 2M, N, Table S3**). A large portion of the motif-carrier mRNAs contains both motifs (**Figure S3C**). Notably, MTAup motifs-carrier mRNAs are translated at a higher rate compared to non-carriers (**Figure S3D**) and are enriched for translation, apoptosis, and polyubiquitination processes (**Figure 2O**). Selective translation of key mRNAs in the core networks was confirmed by polysome fractionation coupled to qRT-PCR (**Figure 2P**). The pyrimidine-rich MTAup#2 motif resembles the well-known 5’TOP motif reported to be regulated by mTORC1^28^. In line with the dependence of selective cap-dependent translation on CDK1 rather than mTORC1 during mitotic stress (**Figure 2E, G**), we demonstrated a significant reduction of the translation of MTAup motif carriers upon CDK1 inhibition (**Figure 2P**). Inhibiting CDK1 further rescued polysome peak, indicative of rescue of global translation (**Figure S3E**), supporting CDK1-dependent blockage of global translation (**Figure 1K**). Overall, these data suggest that core mRNA networks that are potentially functional for MTA sensitivity, are marked by unique sequence motifs to facilitate selective translation in a CDK1-dependent manner.

Given the enrichment of polyubiquitination process among selectively translated mRNAs with the MTAup motifs (**Figure 2O**) and the selective translation of ubiquitin/ubiquitin ligases upon MTAs (**Figure 2P**), we sought to test if disruption of proteostasis could be a novel mechanism of MTA-induced cell death. Patient data analysis revealed a significant positive correlation between protein levels of the proteasome and apoptosis scores in MTA-treated breast cancer patients (**Figure 2Q**). Notably, proteasome score was higher after MTA therapy in responder but not in non-responder patients (**Figure 2R, S3F**). To test the functionality of the disruption of proteostasis in response to MTAs, we inhibited proteasome using MG132^29^ which resulted in almost complete blockage of apoptotic cell death and cap-dependent translation under global translation blockage in mitotically arrested cells (**Figure 2S**). Among the strongly selectively translated ubiquitination-related mRNAs from the core network (**Figure 2O, P**), we showed that silencing RPS27A, encoding a fusion protein of ubiquitin and ribosomal protein S27a, totally blocked cap-dependent translation and apoptosis (**Figure 2T**), mimicking proteasome inhibition (**Figure 2S**). To demonstrate the functionality of the MTAup#2 motif within *RPS27A* 5’UTR in terms of MTA-induced selective translation, we transfected MTA sensitive cells with a vector that contains *RPS27A* 5’UTR upstream of *Renilla* luciferase along with the negative control *HPRT* 5’UTR without the motif, followed by MTA treatment (**Fig. 2U**). Docetaxel treatment significantly increased the translation of *RPS27A* 5’UTR-bearing vector, while translation of *HPRT* 5’UTR-bearing vector with no MTAup motif (negative control) did not change (**Figure 2V**). Collectively, these data suggest that CDK1-dependent activation of selective cap-dependent translation causes proteasome-mediated cell death, serving as a novel mechanism of sensitivity to mitotic stress inducers.

### CDK1-mediated TACC3 phosphorylation is critical for MTA-induced selective cap-dependent translation and mitotic cell death

Given the critical role of CDK1 that we identified in determining MTA sensitivity, we next sought to identify potential clinically relevant CDK1 substrates in mitotically arrested cells. To this end, we extracted the phosphorylated proteins in mitotically arrested HeLa cells from a published study^30^ and further filtered them based on their deregulation in patient samples treated with MTA-containing therapy using another published dataset^17^ (**Figure 3A, Table S4**). Among the top 20 most abundant phosphoproteins in mitotically arrested cancer cells, TACC3 (**Figure 3B, C**) and CENPF phosphorylation were significantly induced by MTAs in responder but not in non-responder patients (**Table S4**). Since CENPF has already been shown to be phosphorylated by CDK1 during mitosis^31^, we focused on TACC3 phosphorylation during mitotic stress. Inhibiting CDK1 using RO-3306, but not AURKA (canonical upstream kinase of TACC3^32^) or mTORC1, reduced TACC3 phosphorylation (**Figure 3D**, **2E**), suggesting that TACC3 could be a CDK1 substrate. Indeed, we showed the binding of TACC3 with CDK1 in mitotic cells (**Figure 3E**) and demonstrated that CDK1 directly phosphorylates TACC3 at S558 in an in vitro kinase assay (**Figure 3F**). Overexpressing phospho-defective S558A TACC3 mutant that includes an shRNA sequence against endogenous TACC3^33^ in sensitive cells reduced MTA-induced mitotic arrest, p-4E-BP1, and c. PARP (**Figure 3G**), suggesting that TACC3 phosphorylation has a critical role in mitotic cell death.

**Figure 3.**
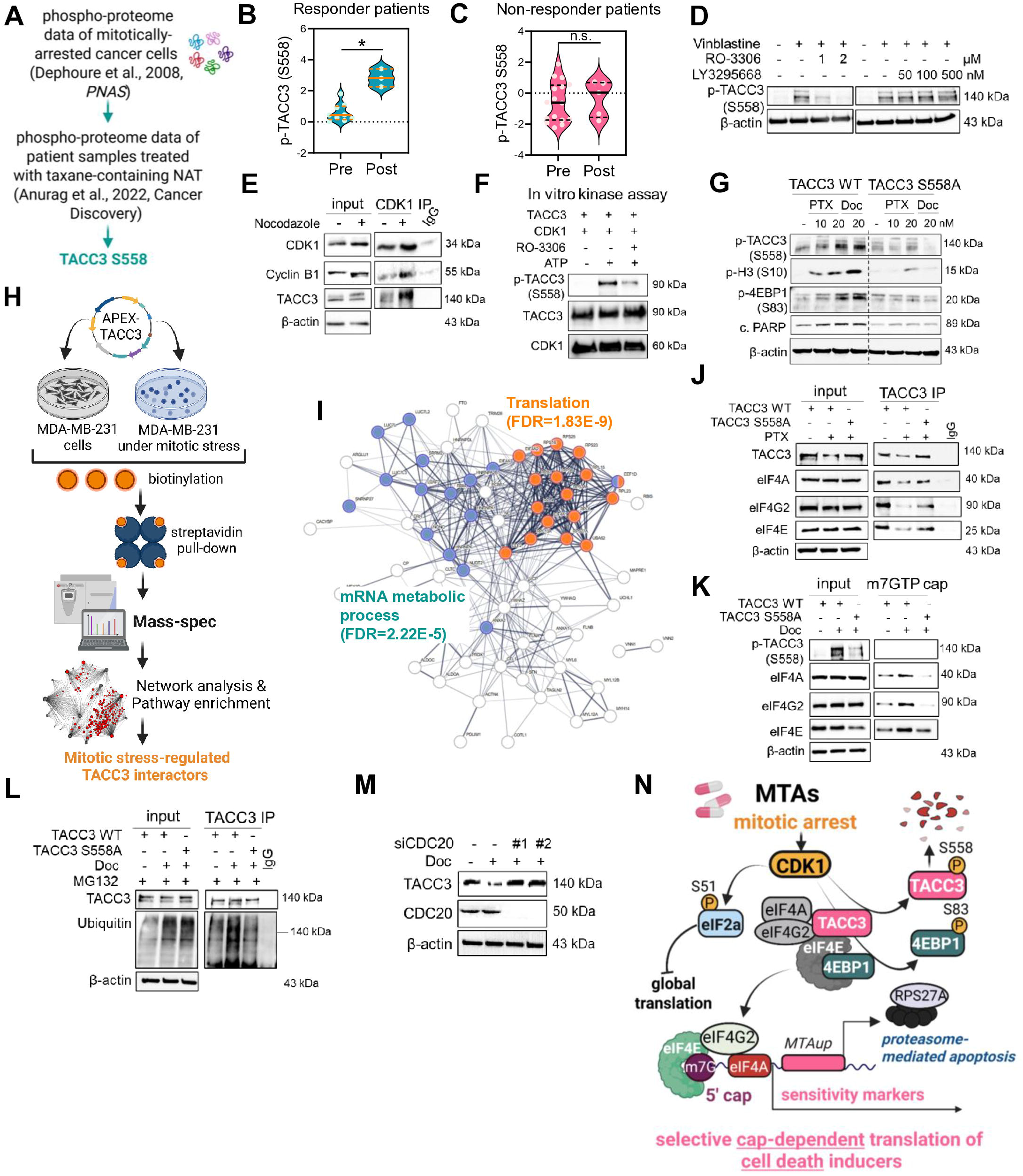
CDK1-mediated TACC3 phosphorylation is critical for MTA-induced selective cap-dependent translation and mitotic cell death. (A) The pipeline of analysis of mass spec data from mitotically arrested HeLa cells and integration with phospho-proteome data from patient samples treated with taxane-containing neo-adjuvant therapy. (B, C) p-TACC3 (S558) levels in responder vs. non-responder patients before and after taxane-containing therapy. (For responders: n=9 pre, n=3 post, for non-responders: n=16 pre, n=5 post). (D) WB of p-TACC3 (S558) in Par cells treated with vinblastine (5 nM) +/− RO-3306 (1 or 2 µM), or the Aurora A inhibitor, LY3295668 (50, 100, or 500 nM). (E) CDK1 IP to show interaction with TACC3 under nocodazole (100 nM). Cyclin B1 is positive control. (F) In vitro kinase assay showing CDK1-mediated S558 phosphorylation of TACC3. (G) WB of markers in wt vs. S558A mut TACC3 expressing cells +/− MTA. (H) Pipeline for TACC3-APEX2 experiment. (I) Pathway enrichment among mitotic stress-regulated TACC3 interactors. (J, K) TACC3 IP (J) and cap pulldown (K) in Par cells overexpressing wt vs. S558A mut TACC3 −/+ MTA. (L) Polyubiquitination assay in the presence of MG132 in Par cells overexpressing wt vs. S558A mut TACC3 −/+ docetaxel (100 nM). (M) WB of TACC3 under docetaxel (50 nM) upon siCDC20. (N) Model of MTA sensitivity. Data represents SD. Two-sided Student’s t-test was used to calculate statistical difference between two groups. ** *P*<0.01; n.s., not significant.

To identify novel and physiologically relevant interactors of TACC3 that might be involved in selective translation upon mitotic stress, we performed proximity ligation assay APEX2 combined with mass spectrometry in MDA-MB-231 cells synchronized in mitosis vs. interphase (**Figure 3H**). We found that among 96 TACC3 interactors in interphase (FC cut-off = 1.5), the interaction of 79 of these proteins with TACC3 was downregulated (FC cut-off = 1.5) in mitotically arrested cells (**Table S5**). Downstream analysis with the mitotic stress-regulated interactors revealed enrichment of translation (FDR=1.83 × 10^−9^) and mRNA metabolic process pathways (FDR=2.22 × 10^−5^) (**Figure 3I**). These data suggest that TACC3 serves as a hub for translation-related proteins, which are released upon CDK1-mediated TACC3 phosphorylation to execute selective translation. Indeed, we identified and validated initiation factors eIF4A, eIF4G2, and eIF4E, as novel TACC3 interactors and showed that their interactions with TACC3 were reduced by MTAs only in wt TACC3, but not in S558A TACC3 mutant-expressing cells (**Figure 3J**). The reduced interaction of wt TACC3 with the initiation factors led to increased cap recruitment of the initiation factors that was reduced in the presence of S558A TACC3 mutant (**Figure 3K**). Notably, we detected a downregulation of wt but not S558A mutant TACC3 protein by MTA (**Figure 3J**) that is potentially driven by increased polyubiquitination (**Figure 3L**). Indeed, silencing CDC20, activator of the mitotic ubiquitin ligase, APC/C^34^, reversed MTA-induced TACC3 downregulation (**Figure 3M**). Collectively, these data suggest that MTA-induced selective cap-dependent translation is mediated by the inhibitory S558 phosphorylation of TACC3 that releases a pool of initiation factors to selectively translate pro-apoptotic mRNAs under MTA-induced mitotic stress (**Figure 3N**).

### TACC3-driven cap-independent translation of m^6^A methylated mRNAs with hnRNPC motif leads to faithful mitosis in MTA tolerant cells

We next sought to determine the mechanisms of MTA tolerance, a poorly characterized reversible precursor state before full-blown resistance. We developed MTA tolerant cells by culturing the sensitive cells in the presence of MTAs for 10 days (**Figure S4A**) as reported before ^35^. Tolerance was reversible as drug withdrawal and re-seeding caused regain of sensitivity (**Figure S4B, C**). Puromycin labeling revealed that similar to sensitive cells under MTA treatment, tolerant cells also had reduced rates of global translation (**Figure S4D**). Interestingly, unlike sensitive cells, blockage of global translation in tolerant cells was followed by increased cap-independent translation (**Figure 4A, B**), suggesting a switch from cap-dependent to cap-independent translation during MTA tolerance. In line with the lack of cap-dependent translation in tolerant cells, mitotic arrest, p-4E-BP1, p-TACC3, and cleaved PARP were lower compared to sensitive ones (**Figure 4C**).

**Figure 4.**
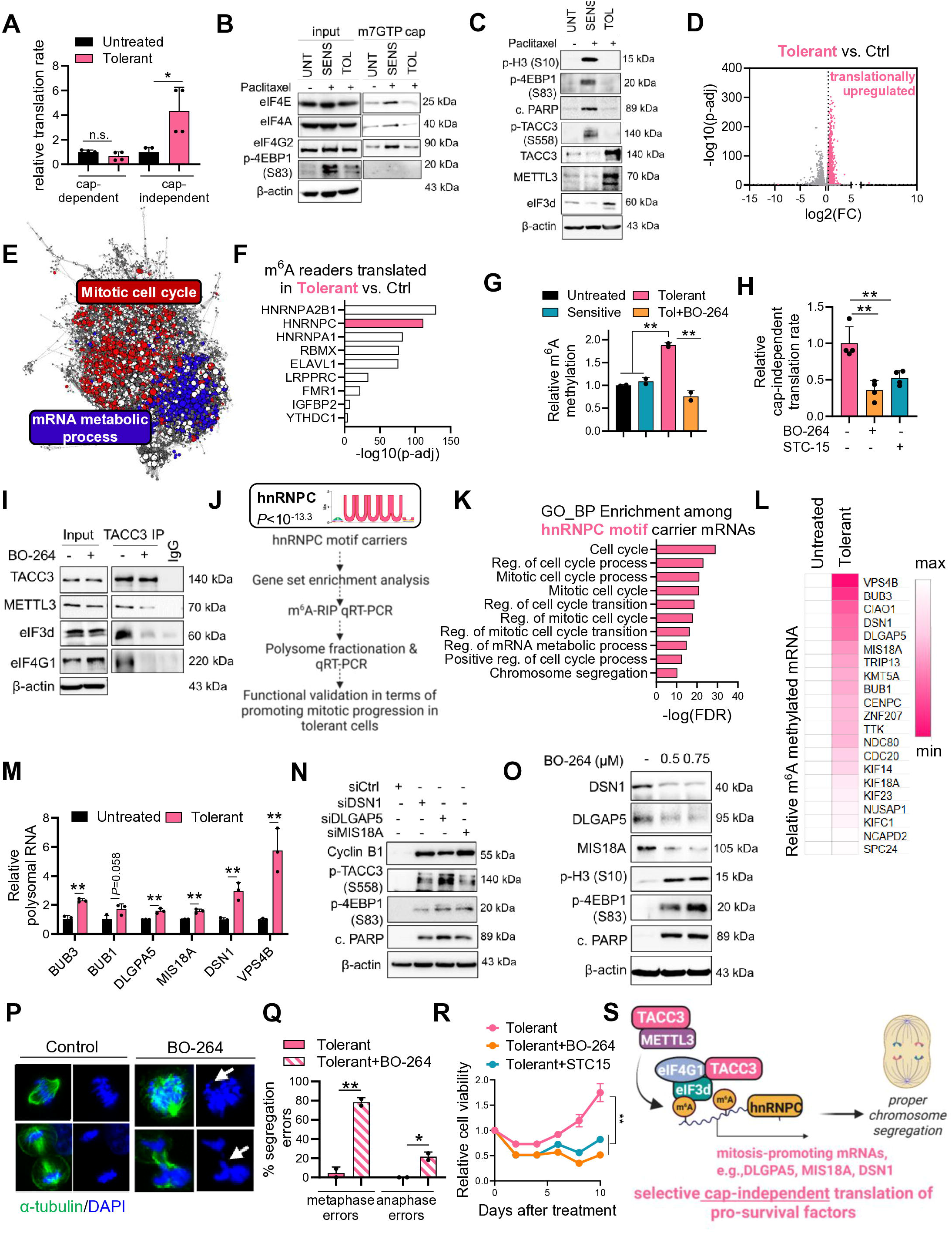
TACC3-driven cap-independent translation of m^6^A methylated mRNAs with hnRNPC motif leads to faithful mitosis in MTA tolerant cells. (A) Cap-dependent and -independent translation rate in untreated vs. tolerant cells (n=4). (B, C) Cap pull-down (B) and WB of sensitivity and potential tolerance mediators (C) in untreated, sensitive, and tolerant cells. (D) Volcano plot of selectively translated mRNAs in tolerant cells determined by Polysome-seq. (E) Network analysis among translated mRNAs in tolerance. (F) Translation of m^6^A readers in tolerant cells. (G) Relative m^6^A methylation upon TACC3 inhibition in tolerant cells (n=2). (H) Cap-independent translation in tolerant cells treated with BO-264 (1 µM) or STC-15 (10 µM) (n=4). (I) TACC3 IP in tolerant cells treated with BO-264 (10 µM) for 4 hrs. (J) Overview of the strategy employed to determine selectively translated mRNAs with hnRNPC motif that are functionally important for mitotic progression in MTA tolerant cells. (K) Pathway enrichment among hnRNPC motif carrier mRNAs selectively translated in tolerant cells. (L) Heatmap of relative m^6^A methylation of hnRNPC motif carrying selectively translated mRNAs related to chromosome segregation in tolerant cells compared to untreated cells. (M) Relative polysomal enrichment of chromosome segregation-related m^6^A methylated mRNAs from K in untreated vs. MTA-tolerant cells (n=3). (N) WB analysis of mitotic arrest and cell death markers in tolerant cells transfected with siRNAs against DSN1, DLGAP5, and MIS18A in the presence of paclitaxel (5 nM) for 24 hours. (O) WB of DSN1, DLGAP5, and MIS18A, and the markers in tolerant cells treated with BO-264 (0.5, 0.75 µM) for 24 hours. (P, Q) IF staining of α-tubulin and DAPI (P) and quantification of segregation errors (Q) in BO-264-treated (500 nM) tolerant cells. (R) Relative cell viability showing inhibition of tolerance by BO-264 or STC-15 (n=3). (S) Model of MTA tolerance. Data represents SD. Two-sided Student’s t-test was used to calculate statistical difference between two groups except R. In R, paired Student’s t-test was used to calculate statistical difference between groups. \**P*<0.05, ** *P*<0.01; n.s., not significant.

To determine the selectively translated mRNAs in tolerant cells that could facilitate survival, we performed polysome fractionation coupled to RNA-seq (**Figure 4D, Table S6**). We found that mRNAs related to mitotic progression and mRNA metabolic processing were selectively translated in tolerant cells (**Figure 4E**). Of note, mRNA metabolic processing was also significantly enriched among mitotic stress-regulated TACC3 interactors (**Figure 3I**). m^6^A methylation is one of the most abundant and conserved modifications in eukaryotic RNAs^13^ that is involved in numerous aspects of mRNA metabolism, including mRNA processing, nuclear export, decay, and translation^36^. Strikingly, we found that among 15 m^6^A readers, 9 of them were significantly highly translated in tolerant cells (**Figure 4F**). Along these lines, expression of the m^6^A writer METTL3 was higher in tolerant cells, together with TACC3 (**Figure 4C**), suggesting that m^6^A marking/reading of mRNAs could be key to their selective translation in tolerance. In line with this, global m^6^A methylation was higher in tolerant cells (**Figure 4G**) together with higher expression of eIF3d (**Figure 4C**), and inhibiting TACC3 with BO-264^37^ reduced m^6^A methylation (**Figure 4G**). Furthermore, using BO-264^37^ or the METTL3 inhibitor STC-15^38^ significantly reduced cap-independent translation (**Figure 4H**). As an underlying mechanism, we showed for the first time that TACC3 interacts with METLL3 along with eIF3d and eIF4G1 in tolerant cells and TACC3 inhibition by BO-264^37^ reduces this interaction (**Figure 4I**).

Since RNA-binding proteins (RBPs) may also function as m^6^A readers to provide further specificity to mRNA translation, we next sought to test if certain RBP motifs are enriched in the UTR region of selectively translated mRNAs in MTA tolerant cells to functionally contribute to m^6^A methylation, selective translation and MTA tolerance (**Figure 4J**). In line with its higher translation (**Figure 4F**), motif for the m^6^A reader hnRNPC, which is also an RBP, was significantly enriched among the 3’UTR of translationally upregulated mRNAs in MTA tolerant cells (**Figure 4J**). Gene sets related to mitotic progression and chromosome segregation, the most crucial step in mitosis, were strongly enriched within the hnRNPC motif containing mRNAs (**Figure 4K**). Furthermore, m^6^A-RIP coupled to qRT-PCR demonstrated that the chromosome segregation-related mRNAs with the hnRNPC motif are heavily m^6^A methylated in MTA tolerant cells (**Figure 4L**), potentially leading to their selective translation (**Figure 4M**). Among those, silencing DSN1, DLGAP5 or MIS18A induced p-4E-BP1 and mitotic cell death in MTA tolerant cells (**Figure 4N, S4E**). Notably, inhibiting TACC3 reduced the levels of DSN1, DLGAP5, and MIS18A, resulting in p-4E-BP1 and mitotic cell death in MTA tolerant cells (**Figure 4O**, **S4F**). Supporting this, we found that TACC3 mRNA is also translated at a higher rate in MTA tolerant cells (**Figure S4G**) that was accompanied by increased m^6^A methylation (**Figure S4H**), potentially resulting in higher mRNA stability (**Figure S4I**). Inhibiting TACC3 increased chromosome segregation errors (**Figure 4P, Q**) and prevented MTA tolerance (**Figure 4R**). Notably, inhibiting METTL3, the m^6^A methylation-inducing partner of TACC3 using STC-15 recapitulated the effects of TACC3 inhibition in preventing MTA tolerance (**Figure 4R**), showing its functional importance in driving MTA tolerance. This suggests that there is a potential feedforward loop between TACC3 and m^6^A methylation that leads to upregulation of TACC3 in MTA tolerance. Overall, we show, for the first time, that MTA tolerance is driven by cap-independent selective translation of mRNAs marked by m^6^A methylation and hnRNPC motif that are key for chromosome segregation, mitotic progression, and cell survival in a TACC3-dependent manner (**Figure 4S**).

### TACC3 is upregulated in MTA resistance, and its inhibition overcomes resistance to MTAs via suppressing global translation and restoring selective cap-dependent translation

We showed that higher TACC3 expression is associated with worse clinical outcome in taxane-treated breast cancer patients (**Figure 5A**). Consistently, TACC3 was upregulated in the acquired MTA-resistant cells (**Figure 5B**) as well as in a panel of MTA resistant PDXs with known in vivo response to MTAs that were generated from treatment naïve or taxane-treated patients (**Figure 5C, S5**). Silencing TACC3 using CRISPR-Cas9 mediated knockout (**Figure 5E**) or inducible shRNAs (**Figure S6A**) in the presence of MTAs reduced cell viability via causing mitotic cell death (**Figure 5D, E**) and decreased the colony formation (**Figure 5F, S6B**). Notably, MTAs induced p-TACC3 S558 upon mitotic arrest (shown by p-H3), along with increased p-4E-BP1 S83 and c. PARP in responder PDX tumors treated with MTAs compared to non-responder tumors (**Figure 5G**), in line with the data from acquired MTA resistant cell lines (**Figure 1F**, **2A**). On the other hand, TACC3 overexpression conferred MTA resistance (**Figure 5H, S6C**). Targeting TACC3 using BO-264^37^ sensitized resistant cells to MTAs (**Figure 5I**). TACC3 inhibition induced mitotic arrest, blocked global translation, and increased p-4E-BP1 (S83) and c.PARP, thus restoring MTA sensitivity (**Figure 5J, K**).

**Figure 5.**
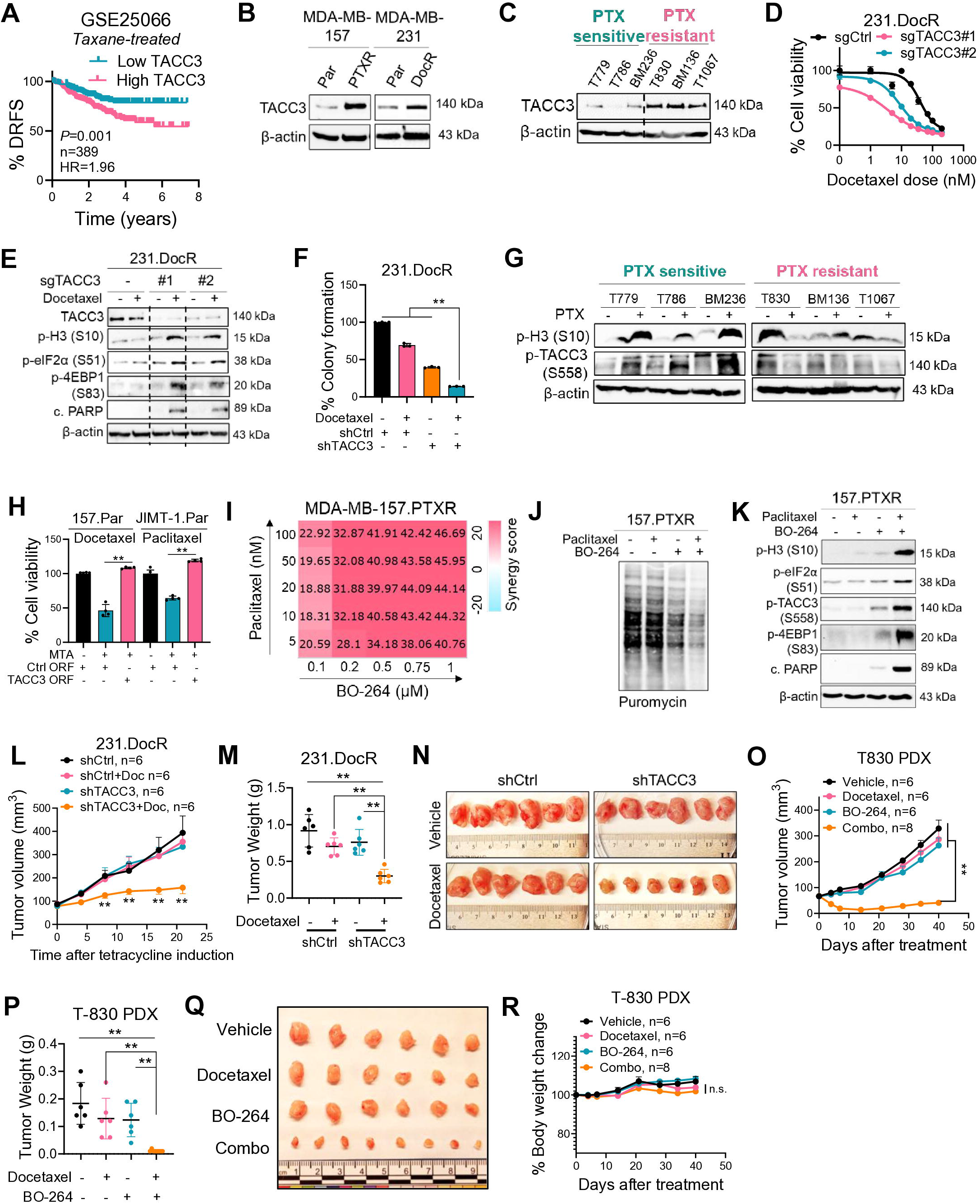
TACC3 is upregulated in MTA resistance and its inhibition overcomes resistance to MTAs via suppressing global translation and restoring selective cap-dependent translation. (A) Kaplan-Meier survival analysis based on TACC3 mRNA expression in taxane-treated breast cancer patients from GSE25066. (B) TACC3 expression in Par vs. MTA-resistant cells. (C) WB of TACC3 in PTX sensitive vs. resistant PDXs. (D) % Cell viability in sgCtrl or sgTACC3-expressing 231.DocR cells treated with increasing doses of docetaxel for 72 hours (n=3). (E) WB of the markers in sgCtrl vs. sgTACC3-expressing 231.DocR cells treated with docetaxel (50 nM) for 24 hrs. (F) Colony formation assay in 231.DocR cells with inducible expression of shTACC3 (n=3). (G) WB of p-H3 (S10) and p-TACC3 (S558) in PTX sensitive vs. resistant PDXs −/+ PTX (10 mg/kg once a week, I.V). (H) % cell viability in TACC3-overexpressing Par cells treated with MTAs (docetaxel: 25 nM, paclitaxel: 5 nM) for 3 days (n=4-6). (I) Synergy map of 157.PTXR cells treated with BO-264 in combination with paclitaxel. (J, K) Puromycin labeling (J) and WB of the markers (K) in 157.PTXR cells treated with paclitaxel (10 nM) and BO-264 (500 nM). (L) Tumor volume of inducible shCtrl vs. shTACC3-expressing 231.DocR xenografts upon inducing the knockdown with tetracycline and treated with 10 mg/kg docetaxel (n=6 different mice). (M) Tumor weights from mice in I (n=6 different tumor). (N) Tumor images from mice in L. (O-Q) Tumor volume (O), tumor weight (P), and representative tumor images (Q) in T-830 PDXs treated with docetaxel (5 mg/kg, twice weekly, i.p.) −/+ BO-264 (75 mg/kg, daily, p.o.) (n=6 different mice for vehicle, docetaxel and BO-264, and n=8 different mice for combo). (R) % Body weight change in mice from O. Data represent mean ± SEM for L and O, and mean ± SD for others. For L, pairwise comparisons between the Combo group and each treatment group were performed using the Wilcoxon rank-sum test. For O, significance was calculated by two-way ANOVA with Dunnett’s multiple comparison test. For others, two-sided Student’s t-test was used to calculate statistical difference between two groups. ** *P*<0.01.

To test the effects of TACC3 depletion on MTA sensitization in vivo, we used the inducible shTACC3 expressing MDA-MB-231.DocR cells (**Figure S6A**). Induction of TACC3 knockdown with tetracycline once tumors are ∼100mm^3^ in combination with docetaxel resulted in significant tumor growth inhibition (**Figure 5L-N, S6D**), overcoming MTA resistance. To test the effects of therapeutic targeting of TACC3 on overcoming MTA resistance, we inhibited TACC3 with BO-264^37^ in combination with docetaxel in T-830 MTA resistant PDX generated from a recurrent primary tumor of a docetaxel-treated patient and that overexpress TACC3 (**Figure 5C**). Combination treatment significantly inhibited tumor growth compared to single agent treatments or vehicle treatment (**Figure 5O-Q**) without any major body weight change (**Figure 5R**) or organ toxicity (**Figure S7A, B**). Overall, these data suggest that TACC3 is upregulated in MTA resistance and its inhibition overcomes MTA resistance in vitro and in vivo by blocking global translation and activating selective cap-dependent translation.

### TACC3 and its downstream translation/apoptosis axis are associated with clinical MTA response/resistance

We showed that higher TACC3 expression is associated with worse clinical outcome in taxane-treated breast cancer patients (**Figure 6A**). Furthermore, we performed in situ staining of TACC3 and markers of translation reprogramming in taxane-treated breast tumors from the Q-CROC-03 trial (**Table S7**) before and after therapy and showed that while p-eIF2a, p-4E-BP1, and c.PARP were induced upon treatment in responder tumors they were not changed in non-responders (**Figure 6B-I**). On the other hand, there was a reduction of TACC3 protein in responders after taxane therapy while in non-responders the TACC3 level was stable (**Figure 6J, K**). Together, our results show that higher TACC3 expression is associated with worse outcome, and TACC3 phosphorylation and downstream effectors correlate with clinical MTA response MTA-treated patients.

**Figure 6.**
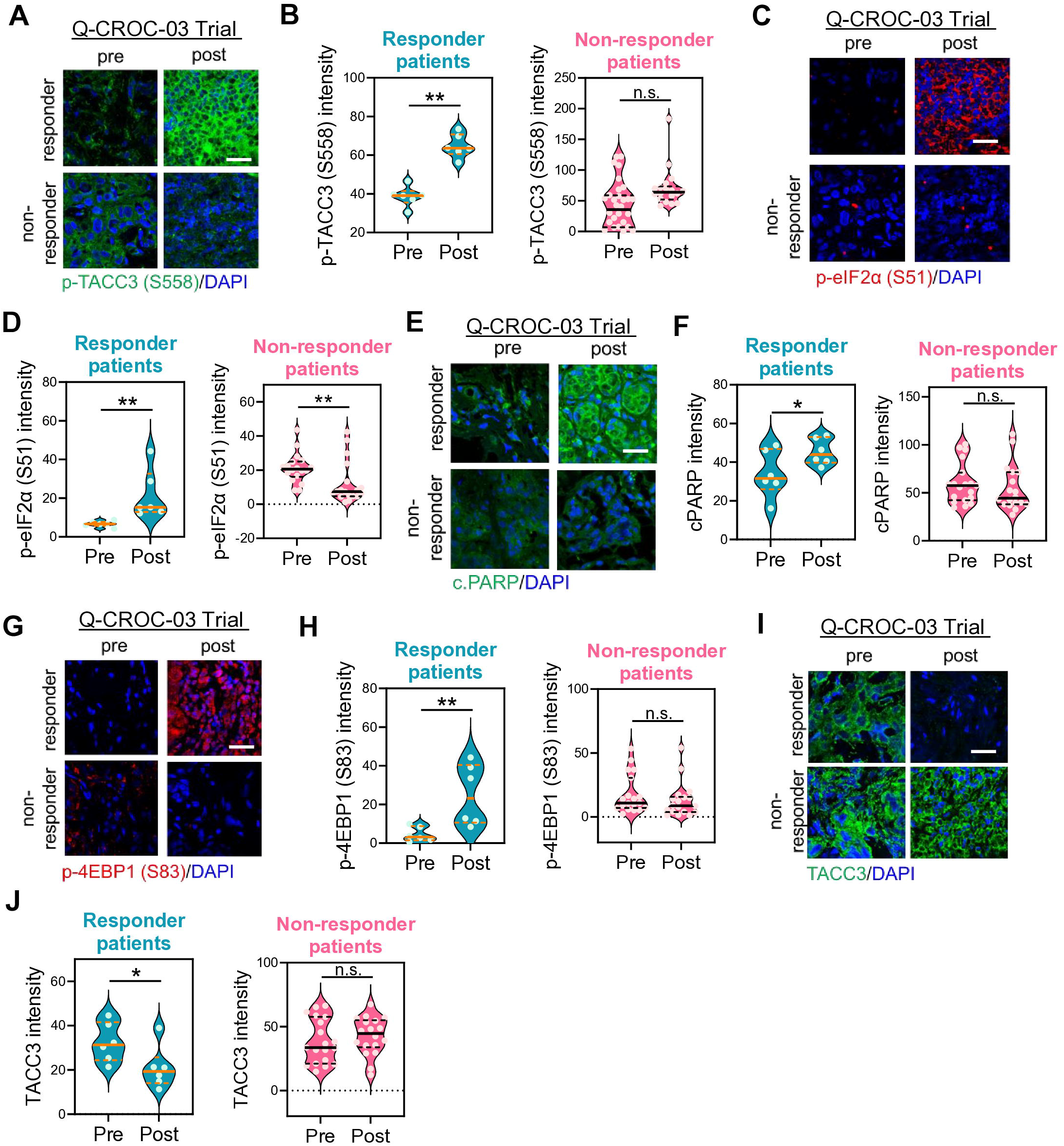
TACC3 and its downstream translation/apoptosis axis are associated with clinical MTA response/resistance. (A-J) In situ IF staining and quantification of the markers in responder vs. non-responder patients before and after taxane-containing therapy from the Q-CROC-03 trial (n=6 areas for pre and post tissues from 3 responder patients and 18-19 areas for pre and post tissues from 10 non-responder patients). Scale bar=50 µM. Significance for survival analysis was calculated by log-rank (Mantel-Cox) test. HR hazard ratio. Two-sided Student’s t-test was used to calculate statistical difference between two groups. \**P*<0.05, ** *P*<0.01; n.s., not significant.

## Discussion

Translation rewiring rapidly alters proteome to allow phenotype switching and adaptation to stress. However, the alternative modes of selective translation and its dynamic crosstalk with epitranscriptomic alterations to determine cell fate during mitotic stress remain largely unknown. Our study uncovers new modes of selective translation governed by specific initiation factors/mRNA sequences and epitranscriptomic mechanisms regulated by TACC3 to orchestrate adaptive responses during MTA-induced mitotic stress. We found that the mitotic stress-induced activation of CDK1 suppresses global translation by phosphorylating eIF2α at S51. On the other hand, it activates selective cap-dependent translation by phosphorylating 4E-BP1 at S83 and TACC3 at S558, leading to TACC3 degradation to promote the recruitment of initiation factors, eIF4A/eIF4E/eIF4G2 to selected mRNAs bearing unique sequence motifs in sensitive cells treated with MTAs. This further disrupts proteostasis, leading to apoptotic cell death (**Figure 3N**). During MTA tolerance, TACC3 is upregulated and interacts with the m^6^A writer METTL3 and the initiation factors eIF3d/eIF4G1 to promote m^6^A methylation and the switch to selective cap-independent translation, leading to mitotic progression and cell survival (**Figure 4S**). The isogenic MTA-resistant cells overexpress TACC3, and TACC3 inhibition overcomes MTA resistance in taxane-resistant PDXs without toxicity in vivo (**Figure 5**). Higher TACC3 correlates with worse clinical outcome in MTA-treated patients (**Figure 5A**).

A few studies have identified transcriptional and epigenetic alterations in cell metabolism, epithelial-to-mesenchymal transition programming, and hijacking of the tumor microenvironment as adaptive responses during cancer cell plasticity^39^. However, this is the first study to report the dynamics of translation reprogramming governed by specific initiation factors/mRNA sequences and epitranscriptomic mechanisms as the key adaptive response to therapy-induced mitotic stress and its contribution to phenotype switching that ultimately dictate therapy response and resistance. The differential modes of selective translation (cap-dependent and independent) and unique translation initiation complexes that we identified in distinct cell states (here: eIF4E/eIF4A/eIF4G2 complex in MTA sensitive and eIF4G1/eIF3d complex in MTA tolerant) lay the foundation for future studies on “translation switching” by epitranscriptomic modifications as a new way of cell adaptation to therapy-induced stress.

The ubiquitin-proteasome system (UPS) is a complex and multifaceted pathway crucial for modulating a vast number of cellular processes, including adaptation to cellular stress and proteostasis, i.e., protein homeostasis^40^ that can either trigger or inhibit apoptosis^41^. Despite having pivotal roles in cancer cell functioning, the mechanisms of proteostasis reprogramming during therapy-induced stress and its crosstalk with other adaptive response pathways are mostly unknown. Our results show, for the first time, that mRNAs of the UPS elements share common sequence motifs that drive their CDK1-mediated selective cap-dependent translation during MTA-induced mitotic stress. We identified the proteasomal *RPS27A*, one of four genes in humans that encode ubiquitin, as a functional mediator of MTA-induced cell death by disrupting proteostasis. *RPS27A* encodes a fusion protein that is post-translationally processed to release free ubiquitin and the mature RPS27a ribosomal protein. In line with this, we showed that RPS27a not only regulates cell death but also selective translation during MTA-induced mitotic stress (**Figure 2T**), likely by facilitating eIF4E/eIF4G2/eIF4A-mediated selective cap-dependent translation. Furthermore, based on our data showing APC/Cdc20-mediated proteasomal degradation of TACC3 upon mitotic stress (**Figure 3M**), it is likely that RPS27a-ubiquitin is incorporated into the APC/C complex that is activated by Cdc20 to facilitate TACC3 degradation.

The precise control of selective translation of functionally linked sets of mRNAs during cellular stress requires both cis-regulatory elements, e.g., m^6^A methylation, and trans-regulatory elements, e.g., RBPs^42^. m^6^A methylation regulates various aspects of RNA biology, including translation, stability, and polyadenylation^13^. Despite having pivotal roles in tumor progression^43^, the potential roles of m^6^A methylation and its modifiers in epitranscriptomic regulation of translation reprogramming during adaptation to therapy (here: tolerance to mitotic stressors) are not known. Furthermore, the potential interplay between cis- and trans-regulatory elements to regulate translation is still not fully deciphered. This is the first study to show the multilayered regulation of mitotic progression during MTA tolerance that involves marking of mRNAs driving chromosome segregation simultaneously with the hnRNPC motif and m^6^A methylation that ultimately increases their selective translation. We identified the TACC3/METTL3 complex as the key targetable driver of epitranscriptomic translation rewiring promoting MTA tolerance. Given that hnRNPC is both an RBP and also an m^6^A reader, it is likely that the presence of hnRNPC motif in tolerance-promoting mRNAs cooperates with the m^6^A methylation by the TACC3-METTL3 complex to effectively recruit initiation complexes.

We identified TACC3 as the coordinator of phenotype switching upon therapy-induced mitotic stress by undergoing multiple layers of regulation, including post-translational modifications, e.g., phosphorylation and ubiquitination (in the sensitive state) and epitranscriptomic modifications, e.g., METTL3-driven m^6^A methylation (in the tolerant state). We deciphered the functions of TACC3 and its unique interactomes in different cell states from sensitivity to tolerance and resistance during MTA-induced mitotic stress. We showed that while TACC3 phosphorylation and degradation by CDK1 in MTA sensitive cells activates selective cap-dependent translation, m^6^A methylation of its mRNA increases stability in tolerance, thus switching the gears towards cap-independent translation of pro-survival factors. The robust multilayered regulation of TACC3 provides evidence for its centrality for adaptive responses and suggests that TACC3 may act as a molecular rheostat, adjusting translation responses to therapy-induced mitotic stress to facilitate cell death or fuel cell survival.

In summary, our study identified the molecular mechanisms of translation/epitranscriptomic reprogramming governed by formation of alternative initiation complexes and altered m^6^A methylation as an adaptive response to mitotic stress. We also identify the key multi-functional adaptor protein, TACC3 as the master orchestrator of translational/epitranscriptomic reprograming and the driver of tolerance/resistance to MTAs, the cornerstone mitotic stress inducer therapy. We showed that targeting TACC3 has therapeutic potential to restore MTA sensitivity, representing a promising therapeutic option for patients with refractory disease.

## Limitations of the study

There are several limitations in this study. First, we showed that mitotic selective cap-dependent translation and cell death is dependent on CDK1. However, we have not demonstrated how CDK1 is activated during MTA-induced mitotic stress. Second, we identified the sequence motifs that potentially mark the mRNAs to be selectively translated in sensitive vs. tolerant cells. We also demonstrated an association between hnRNPC motif and m^6^A methylation for the selectively translated mRNAs. However, whether RNA modifications other than m^6^A methylation could also be critical need further investigation. Third, we showed that TACC3 is upregulated in MTA tolerance potentially via increased m^6^A methylation and stability of its mRNA, despite the lack of the hnRNPC motif. This necessitates further investigation to identify the regulators of m^6^A methylation of TACC3 mRNA during MTA tolerance.

## Resource availability

### Lead contact

Further information and requests for resources and reagents should be directed to and will be fulfilled by the lead contact, Ozgur Sahin.

### Materials availability

Reagents used and generated in this study are available from the lead contact upon request.

### Data and code availability

- All datasets generated in this study are publicly available, and accession numbers are also listed in the key resources table.
- Raw fastq files resulting from the polysome-seq studies are deposited in the National Center for Biotechnology Information (NCBI) Gene Expression Omnibus (GEO: GSE304698).
- The mass spectrometry proteomics data from APEX2 TACC3 experiment have been deposited to the ProteomeXchange Consortium via the PRIDE^44^ partner repository with the dataset identifier PXD067477.

This paper analyzes existing, publicly available datasets. Accession numbers for these datasets are listed in Star Methods.

- This paper does not report original code. Software and algorithms used in this study are listed in key resources table.
- Any additional information required to reanalyze the data reported in this paper is available from the lead contact upon request.

## Acknowledgments

We are thankful to the members of the Ozgur Sahin laboratory for their invaluable discussion and advice. We thank the Translational Science Laboratory and the Flow Cytometry and Cell Sorting Shared Resource of the Medical University of South Carolina. We thank Dr. Stephen Royle (University of Warwick) for providing TACC3 ORF-expressing vector. This work was supported by research funding from the DOD, BCRP (HT9425-26-1-E012 to O. Sahin), and in part from the NIH (R01CA251374 and R01CA267101 to O. Sahin) and SmartState Endowment in Lipidomics and Drug Discovery (O.Sahin), and Hollings Cancer Center, MUSC (Postdoctoral Fellowship to O. Saatci). The core facilities utilized are supported by NIH (C06 RR015455), Hollings Cancer Center Support Grant (P30 CA138313). The Zeiss 880 microscope was funded by a Shared Instrumentation grant (S10 OD018113). The proteomics project was performed in the MUSC Mass Spectrometry Facility and supported, in part, by NIH grants S10 OD028692, P30 CA138313, and P20 GM103542. The Jewish General Hopsital Breast Cancer biobank contributed tissues for PDX generation and is supported by FRQS Réseau Recherche Cancer and the Quebec Breast Cancer Foundation. The Q-CROC-03 clinical trial was supported by Genome Quebec, McGill University and the Jewish General Hospital Foundation. AAM is supported by the Guerrera Family Scientist Award.

## Authors Contributions

**O. Saatci:** Conceptualization, resources, data curation, formal analysis, validation, investigation, visualization, methodology, writing–original draft, writing–review and editing. **A. H. Corchado:** Data curation, investigation, data analysis. **A. Aguilar-Mahecha:** Data curation, investigation, data analysis. **B. Howley:** Data curation, investigation, methodology, writing– review, and editing. **J. Rutherford Bethard:** Data curation, investigation. **Jean-Sebastien:** Statistical analysis. **J. Lafleur**: Data collection. **M. Buchanan**: Methodology and data collection. **E. G. Hill:** Statistical analysis. **L. E. Ball:** Data curation, investigation, writing–review, and editing. **S. Mathew:** Data curation, investigation, methodology, writing–review, and editing. **P. Howe:** Data curation, investigation, methodology, writing–review, and editing. **M. Basik:** Data curation, investigation, writing–review, and editing. **H. Najafabadi**: Data curation, investigation, methodology, writing–review, and editing. **O. Sahin:** Conceptualization, resources, formal analysis, supervision, funding acquisition, validation, visualization, methodology, project administration, writing-review, and editing.

## STAR★Methods

### Key resources table

#### Experimental model and study participant details

##### Human Tissue Samples

Archival paraffin-embedded human breast carcinoma tissue samples from the Q-CROC-03 trial (NCT01276899) ^45^ were obtained from the Jewish General Hospital/ Lady Davis Institute and were coded without any patient identifiers. Patients were recruited at five hospital centers (4 in Montreal, QC, and 1 in Chicago, IL) under a protocol reviewed and approved by the local ethics committees of each institution (Comité d’éthique de la recherche du CHUM, Cook Country Health’s Institutional Review Board Research Ethics Boards of the McGill University Health Center, Comité d’éthique de la recherche de l’HSCM and Jewish General Hospital Research Ethics Office). Human tissues used for the generation of PDX models were obtained from breast cancer patients participating in the Jewish General Hospital Breast cancer biobank (protocol 05-006) approved by the Jewish General Hospital REB. All research was performed in accordance with local and international regulations and guidelines, and all patients provided written informed consent.

##### TNBC PDX xenograft mice tumor model

2-3 mm^3^ of PDX tumor pieces from TNBC PDX T-830 were transplanted near mammary fat pad (MFP) of NSG mice. When the tumors reached 75 mm^3^, mice were randomly distributed to 4 groups and treated with vehicle, docetaxel (5 mg/kg twice weekly, i.p.), BO-264 (75 mg/kg daily, oral gavage) or their combination. Tumor volumes were measured using a caliper, and body weights were also recorded for 40 days. For the PDX panel from MTA-treated breast cancer patients (Lady Davis Institute) (**Figure S5**), mice were randomly distributed to 2 groups when the tumors reached 200 mm^3^ and treated with vehicle or paclitaxel (10 mg/kg once a week, i.v.) for 21 days. Response to PTX was determined by calculating tumor volume change from baseline.

For testing the effects of TACC3 knockdown on MTA sensitization in vivo, 3.5 million MDA-MB-231.DocR.shCtrl or shTACC3 (tetracycline inducible) cells were injected near MFP of the NSG mice. When the tumors reached 100 mm^3^, mice were randomly distributed to 4 groups. All mice received tetracycline from drinking water (1 mg/kg) and half of the mice from each of the shCtrl and shTACC3 mice were treated with vehicle or docetaxel (10 mg/kg twice weekly, i.p.). Tumor volumes were measured using a caliper twice a week, and body weights were also recorded. After 21 days of treatment, mice were sacrificed. All the in vivo studies were carried out in accordance with the Institutional Animal Care and Use Committee of Medical University of South Carolina.

##### Cell lines, drugs, and culture conditions

Human breast cancer cell lines, MDA-MB-231 and MDA-MB-157 were purchased from ATCC (VA, USA), and JIMT-1 cell line was purchased from DSMZ (DE). The CDK1 inhibitor RO-3306, mTORC1 inhibitor RAD001, SAC inhibitor TC Mps1, proteasome inhibitor MG132, MTAs (paclitaxel, docetaxel, and vinblastine) and METTL3 inhibitor STC-15 were purchased from MedChemExpress (NJ, USA). BO-264 was purchased from BioDuro sundia (CN). Cells were cultured in DMEM with 10% FBS, 50 U/mL penicillin/streptomycin, 1% nonessential amino acids (Gibco, MA, USA). MTA-resistant cells (231.DocR, 157.PTXR, and JIMT-1.VnbR) were generated by continuous treatment with the respective drugs for over 6 months. The parental counterparts (Par) were cultured alongside resistant cells without drug treatment. Cells were routinely tested for mycoplasma contamination using MycoAlert detection kit (Lonza, CH) and were authenticated by STR sequencing.

### Method details

#### Drug treatments and transient transfection with overexpression vectors

Treatments with MTAs were done for 24 hours for WB analysis and for 3 days for cell viability. Treatment with CDK1 inhibitor RO-3306, mTORC1 inhibitor RAD001, SAC inhihitor TC Mps1, Aurora A inhibitor LY3295668, proteasome inhibitor MG132 and the CREB inhibitor 666-15 was done at the last 6 hours of MTA treatment. Transfections with overexpression vectors listed in **Table S10** were done in P/S-free growth medium with reduced serum at a concentration of 100 nM using Lipofectamine 2000^TM^ (Invitrogen, MA, USA) as previously described^46^. Cells were transfected with the wt (Addgene, 59356) vs S558A mutant (Addgene, 59357) TACC3 vectors 24 hr before treatment with MTAs.

#### Stable transfections using lentiviral vectors

Inducible lentiviral TACC3 shRNA vector (RHS4696-200764244, ACACAACCTCTTCGAACCT) was obtained from Dharmacon. The sgRNA sequences targeting TACC3 in MDA-MB-157.PTXR and MDA-MB-231.DocR cells are 5’-CAGGCAACGTACCCTCAGCG-3’, and 5’-GACTTGGTGTCACCTCCGAA-3’. sgRNAs were designed and selected based on having high on-target (=high efficacy) and low off-target (=high specificity) activity using the CRISPick tool (Broad Institute). The designed sgRNAs were cloned into human lentiCRISPR v2 vector (Addgene, MA, USA). For lentiviral packaging, HEK293T cells were transfected with shRNAs or sgRNAs and the packaging plasmids, pMD2.G and psPAX2 (Addgene, MA, USA). 48 hours post-transfection, viral particles were harvested and used to transduce MDA-MB-157.PTXR and MDA-MB-231.DocR cells for sgTACC3, and MDA-MB-231.DocR cells for shTACC3 in the presence of 10 ug/ml polybrene. 96 hours post-transduction, stably transfected cells were selected with 2 μg/ml of puromycin for 3-4 days.

#### Whole-transcriptome sequencing (RNA-seq) and data analysis

RNA-seq of polysome-bound and total RNA from MTA sensitive and tolerant cells were performed in duplicates using the Illumina HiSeq 2000 platform at McGill University and Genome Quebec Innovation Centre as described previously ^47^. Raw paired reads were used to quantify transcript abundances using Kallisto v0.46.1 (parameters: –-rf-stranded). Transcript information was obtained from Ensembl (release 111). Transcript-level abundance estimates were collapsed to the gene level using the ‘‘tximport’’ and count matrices for polysome and whole cell counts were created, genes with a mean smaller than 30 counts were removed ^48^. The determination of translation efficiency was performed using DiffRac analysis ^49^.

#### Polysome fractionation

Cells were treated with 100 µg/mL of CHX for 30 min at 37 ℃. Cell lysis was performed in high salt polysome buffer (20 mM Tris-Hcl pH=7, 150 mM NaCl, 5 mM MgCl2, 100 µg/mL of CHX, 1% Triton X-100 and Protease inhibitor). After 10 min incubation on ice, cells were scraped and triturated 5 times through 26G needle. Centrifugation was performed at 20,000g for 10 min. The supernatant was carefully laid on top of the sucrose gradient prepared the day before. Ultracentrifugation at 35,000 rpm for 3 hrs was done at +4. Fractions were separated by a fractionator and 500 µL of fractions were collected. RNA isolation was done using Trizol LS (Thermo Fisher Scientific, MA, USA).

#### In vitro kinase assay

In vitro kinase assay was performed by incubating 0.2 ug CDK1 recombinant protein (BPS Biosciences, CA, USA) with 0.1 ug 4E-BP1 (SinoBiological, CN) or 0.2 ug TACC3 (Origene, MD, USA) or 0.1 ug eIF2α (Enzo Life Sciences, NY, USA) recombinant proteins at 30^0^C for 60 min in kinase buffer (70 mM Tris-HCl, 10 mM MgCl2, 50 mM DTT) with or without 850 uM ATP or 5 uM of the CDK1 inhibitor RO-3306 as the negative control. The reaction was stopped by transferring samples on ice and proteins were denatured by boiling in 1X SDS sample loading buffer for 10 min at 70 °C. Samples were then loaded onto polyacrylamide gel.

#### Luciferase reporter assay

To test the effects of TACC3 inhibition on cap-dependent and independent translation, we used the pcDNA3 RLUC POLIRES FLUC reporter vector (Addgene, 45642). Vectors with *RPS27A* (ENST00000402285) 5’UTR or *HPRT* (ENST00000298556) 5’UTR upstream of the *Renilla* luciferase were synthesized by Vector Builder (IL, USA) (See **Table S10**). 231.Par cells were transfected with 25 ng or 250 ng of the vectors in 96-well or 6-well plates, respectively using Lipofectamine 2000. Transfected cells were treated with 100 nM docetaxel the next day for a total of 20 hrs. Dual luciferase assay (Promega, WI, USA) was performed with the 96-well plate according to manufacturer’s instructions, while RNA isolation and qRT-PCR of *Renilla* luciferase was performed with the 6-well plate. The primer sequences were provided in **Table S9**. Translation rate upon MTA treatment was calculated by taking the ratio of the luciferase signal to mRNA levels of the vectors in each sample followed by normalizing to untreated control as previously reported ^50^.

#### Site-directed mutagenesis

To mutate 4E-BP1 serine 83 residue to alanine, we utilized the Agilent Technologies Quikchange Xl Site-Directed Mutagenesis Kit (NEB, MA, USA) according to manufacturer’s instructions. The primer sequences were: Forward: GGGGTCACCGCACCTTCCAGT, and Reverse: ACTGGAAGGTGCGGTGACCCC.

#### Cap pulldown

MTA-treated sensitive or tolerant cells were lyzed in lysis buffer (50 mM TrisHCl pH=7.0, 150 mM NaCl, 0.2% NP40, 7.5% glycerol, protease and phosphatase inhibitor cocktail), and clarified by centrifugation. 1 mg protein for each condition was incubated with Immobilized γ-Aminophenyl-m7GTP beads (Jena Biosciences, DE) overnight at +4^0^C. Beads were washed and resuspended in 60 µL 1X SDS sample loading buffer, boiled for 10 min at 70 °C, eluted by centrifugation and loaded onto polyacrylamide gel.

#### Western blotting

Protein isolation and Western blotting were performed as previously described ^37,47,51–53^. Briefly, RIPA buffer was used to isolate total protein lysate in the presence of protease and phosphatase inhibitor cocktails, and protein concentrations were measured using the BCA Protein Assay Reagent Kit (Thermo Fisher Scientific, MA, USA). For extracting proteins from collagen-embedded cells, cells were treated with 1.5 mg/ml of collagenase solution for 5 minutes at 37 °C. Equal amounts of protein were separated using 8-10% SDS-PAGE gel. Separated proteins were transferred to PVDF membranes (Bio-Rad, CA, USA) using a Trans-Blot turbo transfer system (Bio-Rad, CA, USA) and incubated with primary antibodies that are listed in **Table S8**. Horseradish peroxidase-conjugated anti-mouse or anti-rabbit antibodies (Cell Signaling Technology, MA, USA) were used as secondary antibodies, and signals were detected by enhanced chemiluminescence (Thermo Fisher Scientific, MA, USA). Images were acquired using Image Lab Software (Biorad, CA, USA) or iBright Software (Invitrogen).

#### Immunoprecipitation

Cells were treated with 50 nM of MTAs for 24 hours or 10 µM of BO-264 for 4 hr. Cells were lyzed in lysis buffer (50 mM TrisHCl pH=7.0, 150 mM NaCl, 0.2% NP40, 7.5% glycerol, protease and phosphatase inhibitor cocktail), and clarified by centrifugation. 1 mg protein for each condition was incubated with antibody-coated Dynabeads Protein G (Invitrogen, MA, USA) overnight at +4^0^C. Beads were washed and resuspended in 60 µL 1X SDS sample loading buffer, boiled for 10 min at 70 °C and loaded onto polyacrylamide gel.

#### APEX2 proximity ligation assay

JIMT-1 cells were transfected with APEX2-TACC3 vector and synchronized for mitosis and interphase using 100 ng/mL nocodazole and double thymidine block (2 mM), respectively. Biotinylation was performed as previously described ^54^. Briefly, cells were treated with 2.5 mM biotin phenol (Iris Biotech, DE) for 1 hr, and 1 mM H_2_O_2_ was added at room temperature (RT) for 2 min. After washing, cells were lysed with RIPA buffer + quenchers (5mM Trolox, 10 mM NaN_3_, and 10mM Sodium Ascorbate). Lysates were sonicated and clarified by centrifugation. Pre-washed Dynabead M-280 Streptavidin beads were incubated with the cell lysate at 4°C for 4 hr. After the incubation, beads were washed and boiled in elution buffer, and samples were loaded onto polyacrylamide gel for immunoblotting the interactors.

#### Mass spectrometry

Proteins were digested directly off the streptavidin magnetic beads. Beads were washed twice with 50 mM ammonium bicarbonate and the enriched proteins were reduced with 45 mM dithiothreitol and alkylated with 100 mM iodoacetamide. Trypsin (100 ng) was added to each sample and incubated overnight at 37°C with shaking at 300 rpm. The digestion was terminated with the addition of 10% trifluoroacetic acid solution. The supernatant containing the tryptic peptides was desalted using ZipTip with 0.6μL C18 resin (Millipore, MA, USA) and dried under vacuum.

Peptides were separated and analyzed on an EASY nLC 1200 System in-line with the Orbitrap Exploris 480 Mass Spectrometer (Thermo Fisher Scientific, MA, USA). Two µg of peptides were pressure loaded onto a C18 reversed phase column (Acclaim PepMap RSLC, 75 µm x 25 cm (2 µm, 100 Å) Thermo Fisher Scientific, MA, USA). The peptides were separated using a gradient of 0-35% B in 120 min (Solvent A: 5% acetonitrile, 0.1% formic acid; Solvent B: 80% acetonitrile, 0.1% formic acid) at a flow rate of 300 nL/min. Spray voltage was set at 2.2 kV and ion transfer tube temperature at 300°C. Mass spectra were acquired in data-dependent mode with a high resolution (60,000) full scan, mass range of m/z 375-1575, with an automatic gain control target value of 300% and a maximum injection time of 25 ms. Monoisotopic peak determination was enabled. The AGC target value for fragment spectra is set at 100%. Cycle time for MS2 scans was 3 s. The resolution was set at 15,000 with a max injection time of 40 ms. An HCD collision energy of 33% was used for peptide fragmentation. Dynamic exclusion of precursors was set to 20s.

The raw files were searched in Max Quant v2.4.2.0 (Max Planck Institute, DE) against the reviewed human protein database (20,433 entries downloaded from Uniprot in February 2024). The search parameters allowed for two missed cleavages, fragment mass tolerances of 20 ppm, and a minimum of seven amino acids per peptide. Carbamidomethylation of cysteine was set as a fixed modification. Methionine oxidation and acetylation of protein N-termini were set as variable modifications. The false discovery rates for peptide spectral matches and protein identification were set to 1% using a reversed database decoy strategy. The search results were further analyzed and filtered in Perseus v. 1.6.15.0 (Max Planck Institute, DE). Reversed database matches and potential contaminants including streptavidin were removed. Only proteins with >2 peptides were retained. The LFQ protein intensities were log2 transformed. To identify proteins with changes in relative abundance between conditions, a log2 intensity cut-off of 1 was selected.

#### Immunofluorescence staining

Cell fixation was performed with methanol at −20 ℃ for 10 min or 4% paraformaldehyde for 15 min, permeabilized and blocked in 3% BSA-PBS. Cells were incubated in the primary antibodies and secondary Alexa Fluor 647 or 488-labeled antibodies (**Table S8**) for 2 hours at room temperature. Cells were also counterstained with DAPI for 5 min. In situ staining of nascent protein translation was performed using the Click-iT™ Plus OPP Alexa Fluor™ 488 Protein Synthesis Assay Kit (Invitrogen, MA, USA) by following manufacturer’s instructions. Images were acquired using ZEN Black LSM 880 airyscan (Carl Zeiss, DE). Images were analyzed using the ImageJ (Fiji) software (NIH, MD, USA).

#### Immunofluorescence tumor staining

The human tumors from the Q-CROC-03 trial ^45^ were first heated in 60^0^C oven for 1 hour. Deparaffinization was done in Xylene for 15 min two times, followed by washing in 100% and 70% ethanol, sequentially. Antigen retrieval was done in Tris-EDTA (pH=8) buffer for 40 min, and slides were incubated with antibodies against TACC3, p-TACC3 (S558), p-eIF2a (S51), p-4E-BP1 (S83), c.PARP and p-Histone H3 (S10) as listed in **Table S8** at a 1: 50 dilution in 5% BSA-PBS for overnight at +4^0^C, followed by 2 hrs incubation with secondary antibodies at 1:100 dilution at 37^0^C. DAPI solution was prepared in 5% BSA-PBS at a dilution 1:1000 was used for nuclear counterstaining for 5 min at room temperature. Cover slips were mounted with ProLong™ Glass Antifade Mountant (Thermo Fisher Scientific, MA, USA). Imaging was done using Zeiss LSM880 NLO with Airscan Confocal Microscopy and quantification of staining intensity was done using Fiji (ImageJ) software (NIH, MD, USA).

#### Quantitative RT-PCR analysis

Total RNA was extracted from cultured cells using Quick-RNA Miniprep Kit (Zymo, CA, USA) or from polysomes using Trizol LS (ThermoScientific, MA, USA) and cDNAs were generated using RevertAid RT Reverse Transcription Kit (Life Technologies, MA, USA). qRT-PCR analysis was performed with gene-specific primers using LightCycler 480 SYBR Green I Master kit (Roche, CH). *HPRT1* and *ACTB* were used as housekeeping genes. The average Ct value was calculated from triplicates of each sample, and the relative mRNA expression was determined. Sequences of the qRT-PCR primers are listed in **Table S9.**

#### Annexin V/PI staining

Annexin V/PI staining was performed as previously described ^37,55^. Briefly, counted cells for each condition were washed with PBS and incubated with 2 µL of FITC-conjugated Annexin V and PI for 30 min. Data collection was done with BD FACS Diva software (BD, NJ, USA), and analysis of late apoptotic cells was performed by calculating the Annexin V and PI-positive cell percentages using De Novo FCS Express software (CA, USA).

#### Bioinformatics analysis

The microarray data sets, GSE25066 and GSE18728 were downloaded from the GEO database. The proteomic and RNA-seq data from neo-adjuvant MTA-containing therapy-treated breast cancer patient samples were from ^17^. Apoptosis and proteasome scores were calculated by summing up the z-scores of apoptosis and proteasome-related genes retrieved from MSigDB ^56^ for each patient. Network analysis was done using Network Analyst and pathway enrichment was done using String database ^57^. De novo motif discovery and RBP enrichment analysis with the Polysome-seq data was done using MEME software ^58^. Survival curves were generated based on median separation using Kaplan-Meier method, and significance between groups was calculated by Log-rank test. For correlation analysis, Pearson correlation coefficients were calculated. Experiments were repeated two to three times independently with similar results.

#### Quantification and statistical analysis

All the results are represented as mean ± standard deviation (SD) or mean ± standard error of the mean (SEM), as indicated in the figure legends. The flowcharts and schemes were drawn using BioRender (https://app.biorender.com/, Agreement Number: DZ26UFS116). Differences were assessed using the two-tailed unpaired or paired Student’s t-test as indicated in the legends. The significance for survival analysis was done using Log-rank test. In vivo tumor volume data was analyzed using longitudinal linear mixed-effects (LME) models, with the natural logarithm of tumor volume as the outcome variable unless otherwise stated in the legends. Predictor variables included the natural logarithm of baseline tumor volume, time (treated as a categorical variable), treatment group, and their interaction. Tumor weights at the end of the experiment were summarized by treatment group using means, standard errors, and 95% confidence intervals. Pairwise comparisons between the Combo group and each treatment group were performed using the Wilcoxon rank-sum test. *P*-values were adjusted for multiple comparisons using Holm’s method for both tumor volume and tumor weight analyses. Statistical analyses were conducted using GraphPad Prism Software and R version 4.3.0 using the packages nlme (v3.1-164), emmeans (v1.10.4), survival (v3.7-0), and survminer (v0.4.9). The differences were considered statistically significant if *P* < 0.05 and indicated by * or *P* < 0.01 and indicated by ** unless otherwise stated as in the legends.

