## Supplemental Information for "TACC3-driven translation reprogramming dictates susceptibility or tolerance to mitotic stress"

### Supplemental Figures

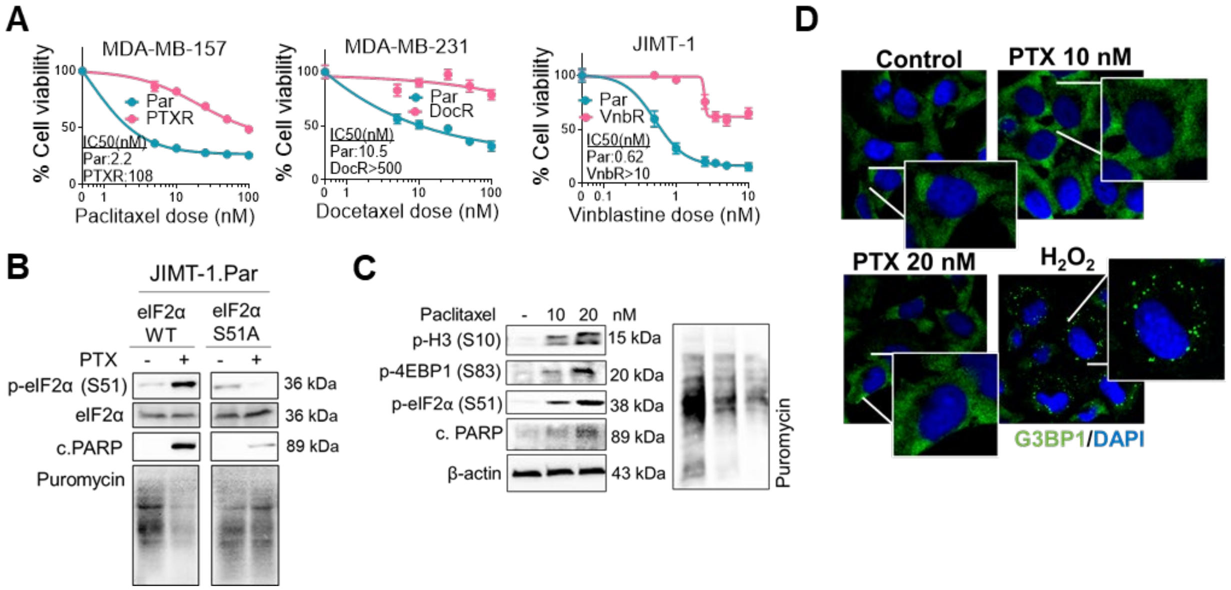


**Figure S1. Dose response curves and IC50s of acquired MTA resistant cell lines vs. parental cell lines and the lack of stress granule formation and mitotic global translation blockage upon MTAs. Related to Figure 1.** (A) Dose response of Par vs. MTA-resistant cell lines after 3 days of treatment of MTAs (n=3-6). (B) WB of p-eIF2α (S51), eIF2α, and c. PARP in JIMT-1.Par cells overexpressing WT vs. S51A eIF2α and treated with paclitaxel (10 nM) for 24 hrs. (C) WB of the markers and puromycin labeling in 231.Par cells treated with paclitaxel (10 or 20 nM) for 24 hrs. (D) G3BP1 staining as the stress granule marker in cells from C. $H_2O_2$ was used as a positive control to induce stress granules.

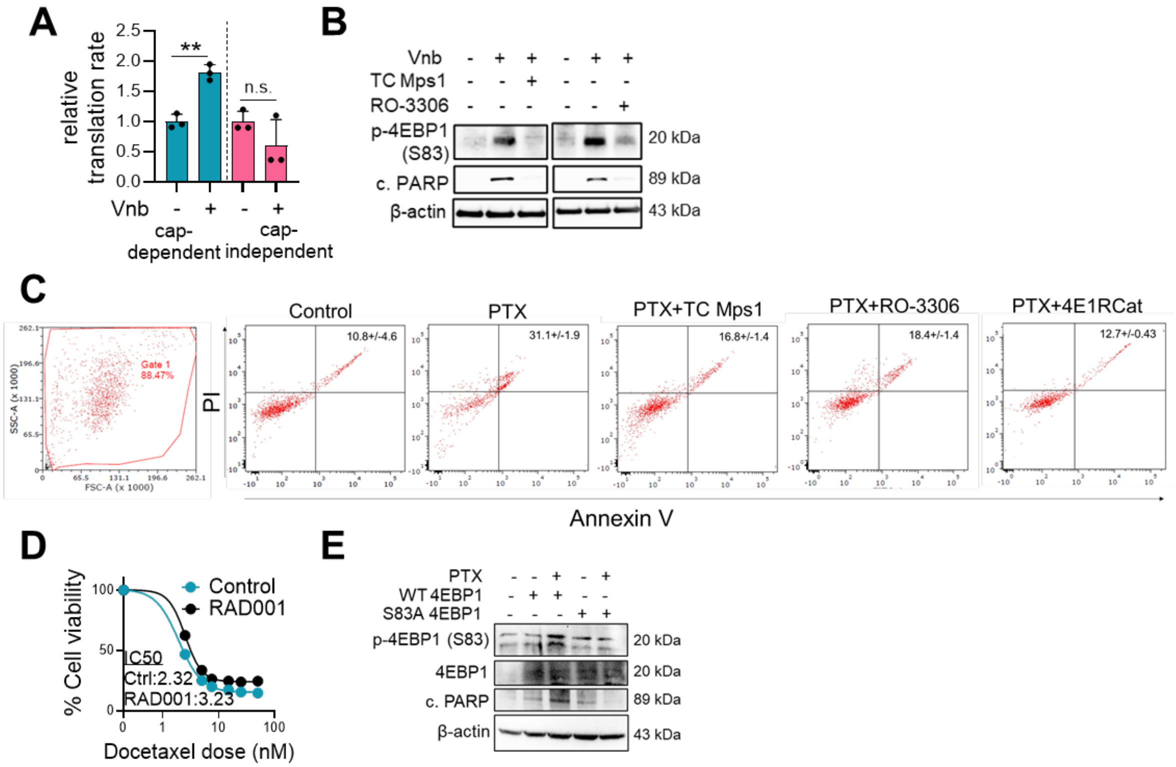


**Figure S2. CDK1-driven 4EBP1 phosphorylation is critical for MTA-induced cell death while mTOR activation is dispensable in MTA sensitivity. Related to Figure 2.** (A) Relative cap-dependent vs. -independent translation rate using luciferase reporter in JIMT-1.Par cells treated with vinblastine for 24 hours (n=3). (B) WB of p-4EBP1 and c.PARP in Par cells treated with vinblastine with or without CDK1 inhibitor RO-3306. (C) Gating strategy and dot plots of cells stained with Annexin V/PI after 24 hours treatment with PTX (50 nM) +/- TC Mps1, RO-3306 (5 μM), or the cap complex inhibitor 4EGI (50 μM) (n=3). (D) % cell viability in JIMT-1.Par cells treated with RAD001 and docetaxel for 3 days (n=4-6). (E) WB of p-4EBP1 (S83), 4EBP1, and c. PARP in Par cells expressing wt vs. S83A mutant 4EBP1 and treated with MTAs.

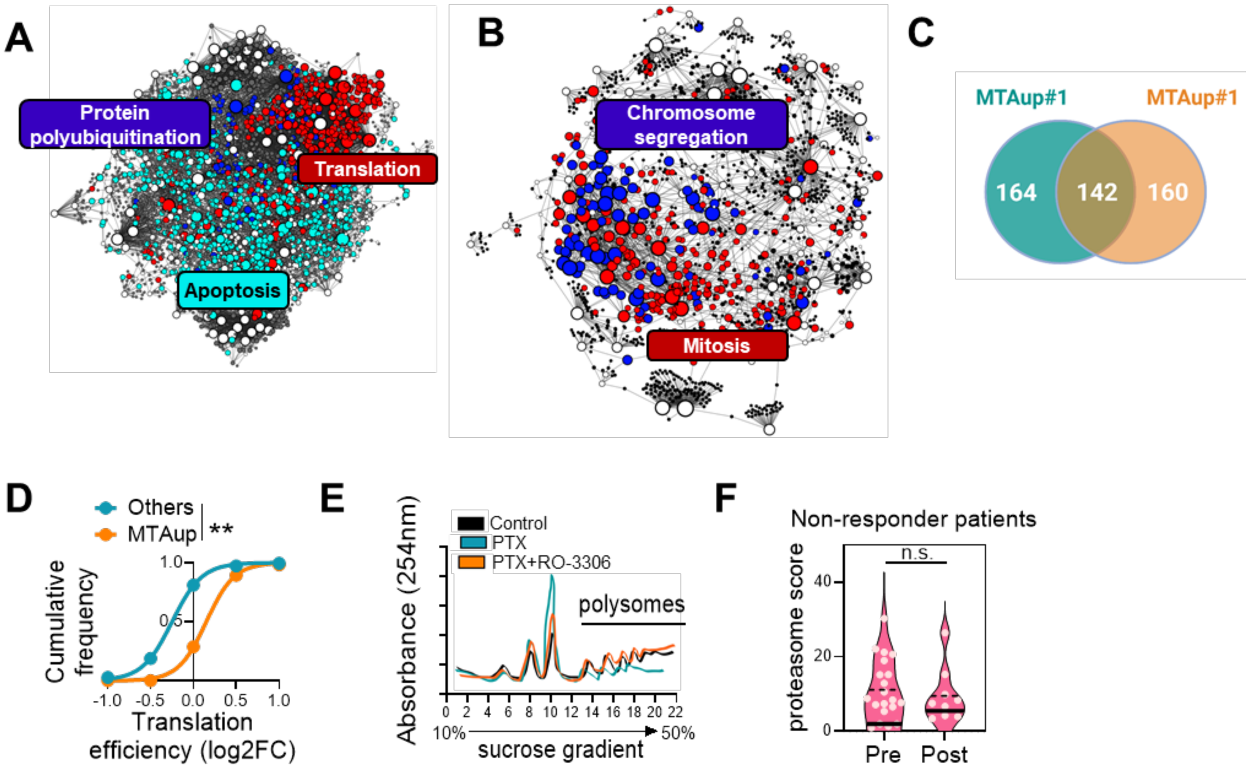


**Figure S3. Network analysis among differentially translated MTA-sensitivity markers, and the role of MTAup motif and CDK1 in translation reprogramming. Related to Figure 2.** (A) Network analysis among translationally upregulated mRNAs upon MTA treatment showing enrichment of translation, apoptosis, and protein ubiquitination. (B) Network analysis among translationally downregulated mRNAs upon MTA treatment showing enrichment of mitosis and chromosome segregation processes. (C) Venn diagram showing the number of mRNAs selectively translated in MTA sensitive cells and bearing MTAup#1 and #2 motifs. (D) Cumulative distribution of translation efficiency of MTAup motif-carrier vs. other mRNAs. (E) Polysome profiles of 231.Par cells treated with PTX with or without the CDK1 inhibitor RO-3306 for 24 hours. (F) Protein levels of the proteasome score in non-responder breast cancer patients before and after taxane-based therapy (n=34 pre, n=12 post).

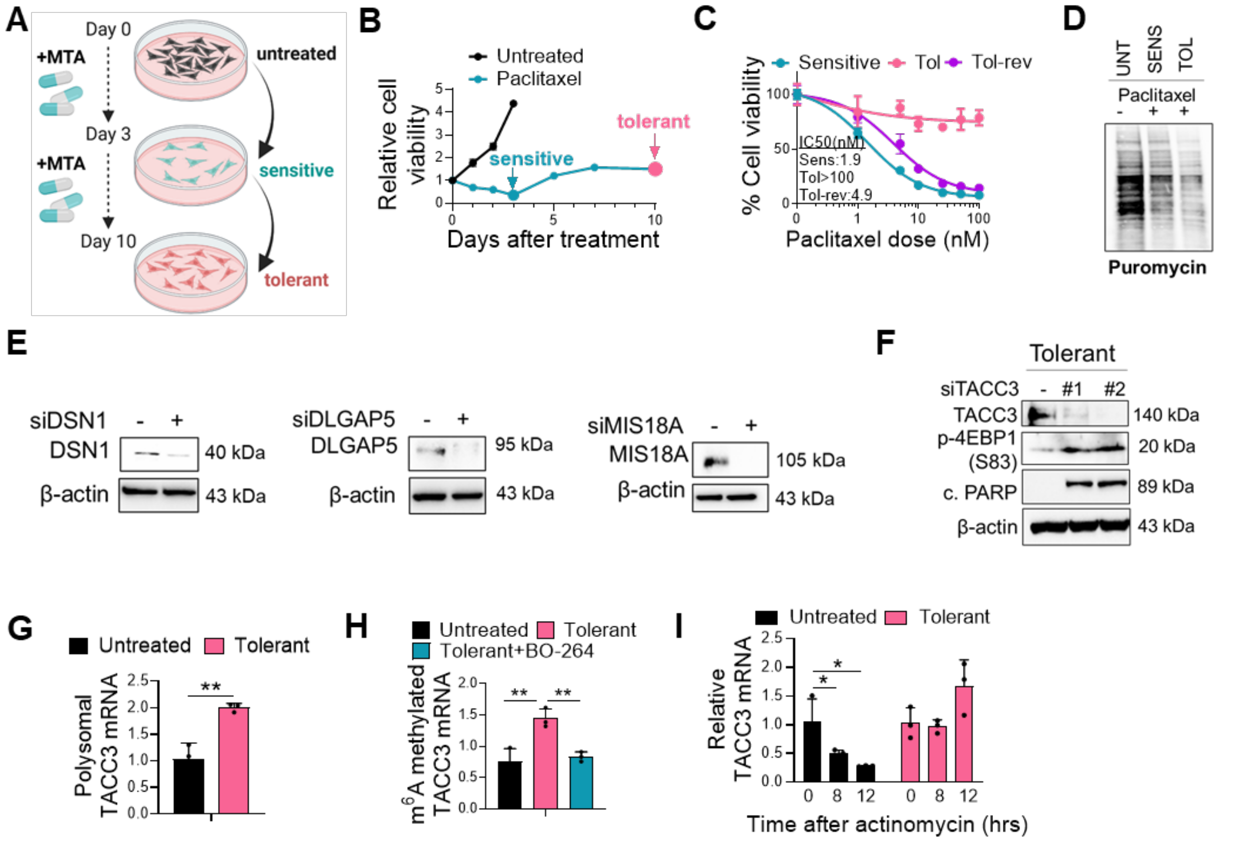


**Figure S4. Generation of MTA tolerance, regulation of TACC3 mRNA stability by m$^6$A methylation and the role of TACC3 in mediating MTA tolerance. Related to Figure 4.** (A) Generation of MTA tolerant cells. (B) Cell viability during tolerance generation (n=4). (C) Dose-response in sensitive vs. tolerant vs. reversed tolerant MDA-MB-157 cells (n=3-6). (D) Puromycin labeling in untreated vs. sensitive vs. tolerant cells. (E) WB validation of DSN1, DLGAP5 and MIS18A knockdown in siRNA-transfected tolerant cells. (F) WB of the markers in tolerant cells transfected with siTACC3 for 24 hours. (G) Polysomal TACC3 mRNA in untreated vs. tolerant cells (n=3). (H) Fold enrichment of m$^6$A binding to TACC3 mRNA (n=3). (I) Relative TACC3 mRNA in untreated vs. MTA-tolerant cells treated with 5 µg/mL actinomycin (n=3).

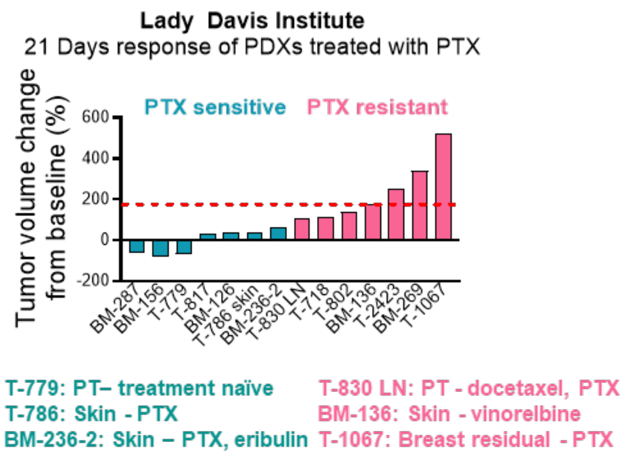


**Figure S5. In vivo taxane response of treatment naïve or taxane-treated breast cancer PDXs. Related to Figure 5.** The in vivo paclitaxel response of a panel of breast cancer PDXs that were generated from treatment naïve or taxane-treated patients with known clinical response.

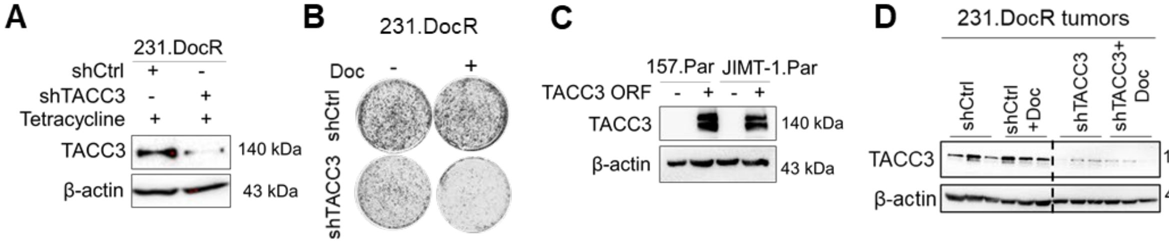


**Figure S6. Inducible TACC3 knockdown in MDA-MB-231.DocR cells and tumors and validation of TACC3 overexpression in Par cells. Related to Figure 5.** (A) WB of TACC3 in shTACC3-expressing MDA-MB-231.DocR cells induced with tetracycline for 3 days. (B) Colony formation assay in 231.DocR cells with inducible expression of shTACC3. (C) WB of TACC3 in TACC3 ORF-expressing 157.Par and JIMT-1.Par cells. (D) WB of TACC3 in shTACC3-expressing MDA-MB-231.DocR tumors induced with tetracycline for 4 weeks from Fig. 5L.

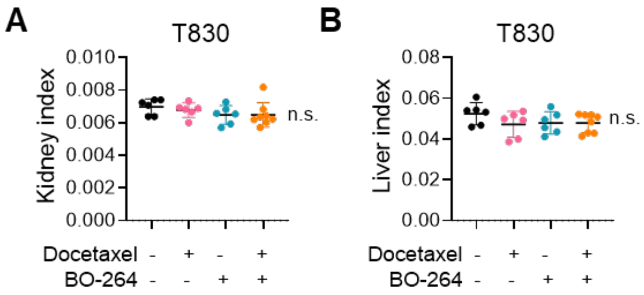


**Figure S7. Potential organ toxicity of the combination of BO-264 and docetaxel in MTA resistant PDXs. Related to Figure 5. A, B.** Kidney (A) and liver (B) indices of mice bearing T-830 PDXs and treated with docetaxel (5 mg/kg, twice weekly, i.p.) -/+ BO-264 (75 mg/kg, daily, p.o.) (n=6 different mice for vehicle, docetaxel and BO-264, and n=8 different mice for combo).

### Supplemental Tables

**Table S1.** mRNAs found in the sub-networks of downregulated mRNAs in MTA-treated patients. Related to Figure 1.

| rRNA metabolic process | tRNA metabolic process | Translation | Mitotic cell cycle-1 | Mitotic cell cycle-2 Poc1a | Sister chromatid segregation |
|---|---|---|---|---|---|
| **DDX49** | GARS | EEF1E1 | BRCA1 | BIRC5 | SEH1L |
| **NOP9** | TARS | GARS | CDK1 | NCAPG2 | SGO2A |
| **RIOK1** | EXOSC7 | TARS | RAD50 | SPDL1 | KNTC1 |
| **BOP1** | IARS2 | MRPS23 | HUS1 | PSRC1 | BUB1B |
| **PRKDC** | NAT10 | IARS2 | TRIP13 | SMC2 | SMC4 |
| **BYSL** | THUMPD3 | MRPL9 | TASOR | ZWINT | KIF4 |
| **EXOSC7** | LARS2 | MRPS15 | FBXL6 | CKAP5 | CIT |
| **WDR12** | DARS2 | AIMP2 | PRKDC | NUF2 | ESPL1 |
| **NIFK** | POLR2L | EIF4EBP1 | MCM4 | NCAPG | NUSAP1 |
| **NAT10** | POLR3K | MRPL3 | MSH2 | DCTN6 | CCNB1 |
| **NOL6** | RPP38 | LARS2 | SEH1L | DNA2 | BIRC5 |
| **UTP11** | ANKRD16 | DARS2 | SGO2A | KNSTRN | NCAPG2 |
| **ERI1** | POP1 | MRPL12 | DTL | CENPA | SPDL1 |
| **EXOSC4** | RPP25 | EFL1 | SAPCD2 | TCF19 | PSRC1 |
| **RPP38** | YRDC | SLBP | PCNA | NABP2 | SMC2 |
| **RRP1** | YARS | MRPL42 | GPSM2 | PRC1 | ZWINT |
| **POLR1A** | TSEN34 | ABCE1 | JTB | WAC | NUF2 |
| **RPP25** | POP7 | RPL10 | CKS1B | KIF14 | NCAPG |
| **UTP25** | PUS7 | GUF1 | STIL | RNF2 | KNSTRN |
| **POLR1B** | ELP4 | TUFM | CDK5 | ANKLE2 | PRC1 |
| **POP7** | TYW5 | YARS | KNTC1 | PDS5A | KIF14 |
| **NHP2** | WDR4 | CKAP5 | WEE1 | KIF20A | PDS5A |
| **RPP40** | TARBP1 | MRPL53 | EIF4EBP1 | CIT | |
| | FARSB | MRPL39 | CCNB2 | ESPL1 | |
| | RPP40 | MRPS17 | BUB1B | NUSAP1 | |
| | | MRPL24 | MZT1 | TUBB3 | |
| | | MRPL15 | DLGAP5 | CCNB1 | |
| | | MRPL17 | TUBB4B | | |
| | | MRPS14 | TUBGCP4 | | |
| | | FARSB | SMC4 | | |
| | | MRPL13 | NUP88 | | |
| | | PDF | KIF4 | | |
| | | | CDC25C | | |
| | | | MCM2 | | |

**Table S2.** List of selectively translated mRNAs in MTA sensitive cells. See the attached Excel Supplementary Table. Related to Figure 2.

**Table S3.** List of selectively translated mRNAs with MTAup#1 and MTAup #2 motifs. Related to Figure 2. See the attached Excel Supplementary Table.

**Table S4.** Top 20 most abundant phosphoproteins in mitotically arrested HeLa cells. Related to Figure 3. The clinically supported proteins are highlighted in red.

| | **Mitotic phosphorylation score** | **Filtering based on association with clinical MTA response*** |
|---|---|---|
| **CENPF** | 275 | + |
| **DLG7** | 197 | - |
| **NUSAP1** | 195 | - |
| **PBK** | 167 | - |
| **ARHGAP11A** | 149 | - |
| **CALD1** | 109 | - |
| **JUN** | 106 | - |
| **CTNND1** | 101 | - |
| **HIST1H2AB** | 100 | - |
| **KIAA1949** | 96 | - |
| **CDC27** | 92 | - |
| **TEX10** | 88 | - |
| **UBAP2** | 88 | - |
| **SEC22L2** | 81 | - |
| **TACC3** | 81 | + |
| **BRRN1** | 79 | - |
| **TTK** | 75 | - |
| **PKMYT1** | 72 | - |
| **PSRC1** | 72 | - |
| **CGI-115** | 71 | - |

*** + represents proteins whose phosphorylation is increased upon MTA treatment in responder patients while it does not change in non-responders.**

**Table S5.** List of mitotic stress-regulated TACC3 interactors. Related to Figure 3. See the attached Excel Supplementary Table.

**Table S6.** List of selectively translated mRNAs in MTA tolerant cells. Related to Figure 4. See the attached Excel Supplementary Table.

**Table S7.** Clinico-pharmacological characteristic of the patients from the Q-CROC-03 trial. Related to Figure 6. See the attached Excel Supplementary Table.

**Table S8.** List of antibodies and their dilutions used in WB, IP, and IF. Related to Figures 1-5.

| Antibody | WB dilution | IP dilution | IF dilution |
|---|---|---|---|
| TACC3 | 1:1000 | 1:10 | 1:400 |
| p-Histone H3 | 1:1000 | | 1:50 |
| Puromycin | 1:1000 | | |
| p-eIF2α (S51), CST3597 | 1:1000 | | |
| p-eIF2α (S51), PT68023-1-Ig | | | 1:50 |
| p-4EBP1 (S83) | 1:1000 | | |
| β-actin | 1:10000 | | |
| p-TACC3 (S558), Thermo PA5-105244 | 1:1000 | | |
| p-TACC3 (S558), CST8842 | 1:1000 | | 1:50 |
| c. PARP, CST9541 | 1:1000 | | |
| c. PARP, CST32563 | | | 1:50 |
| eIF3d | 1:1000 | | |
| eIF4G1 | 1:1000 | | |
| eIF4A | 1:1000 | | |
| eIF4E | 1:1000 | | |
| eIF4G2 | 1:1000 | | |
| METTL3 | 1:1000 | | |
| mTOR | 1:1000 | | |
| CDC20 | 1:1000 | | |
| Cyclin B1 | 1:1000 | | |
| DLGAP5 | 1:1000 | | |
| DSN1 | 1:1000 | | |
| MIS18A | 1:1000 | | |
| Ubiquitin | 1:2000 | | |
| HRP-linked anti-mouse IgG | 1:10000 | | |
| HRP-linked Anti-rabbit IgG | 1:10000 | | |

| | | | |
|---|---|---|---|
| Alexa Fluor 488 anti-mouse | | | 1:100 |
| Alexa Fluor 488 anti-rabbit | | | 1:100 |
| Alexa Fluor 647 anti-mouse | | | 1:100 |
| Alexa Fluor 647 anti-rabbit | | | 1:100 |

**Table S9.** Primer sequences. Related to Figures 2, 4.

| **Primer** | **Sequence** |
|---|---|
| TACC3 forward primer | GTCTGTCTGTCCTGTCTGATTC |
| TACC3 reverse primer | GACAGTGGAGCAGAAGACTAAA |
| NDC80 forward primer | CTGACACAAAGTTTGAAGAAGAGG |
| NDC80 reverse primer | TAAGGCTGCCACAATGTGAGGC |
| DLGAP5 forward primer | AAGTGGGTCGTTATAGACCTGA |
| DLGAP5 reverse primer | TGCTCGAACATCACTCTCGTTAT |
| SPC24 forward primer | GGGATTATGAGTGTGAGCCAGG |
| SPC24 reverse primer | ACTCCAGAGGTAGTCGCTGATG |
| NUSAP1 forward primer | CTGACCAAGACTCCAGCCAGAA |
| NUSAP1 reverse primer | GAGTCTGCGTTGCCTCAGTTGT |
| VPS4B forward primer | GGTTCTGGATTCTGCCATTAGGC |
| VPS4B reverse primer | CCGAAAGTCTGCTTCCGTGAGA |
| KIFC1 forward primer | CCTCACTACAGTGCCACAGACA |
| KIFC1 reverse primer | GAACAGCAGGAACTGGCTTCTG |
| MIS18A forward primer | TGCTTCGCTGTGTTTCCTGT |
| MIS18A reverse primer | TGACACAATTTGCTTTTCAGAGGAC |
| BUB3 forward primer | GTGTGGGACTTACGGAACATGG |
| BUB3 reverse primer | GGGTCCAAATACTCAACTGCCAC |
| ZNF207 forward primer | ATGATGCCACCTGGACCAGGAA |
| ZNF207 reverse primer | GCTGAAACAGCCTGTGCTTGAG |
| TTK forward primer | CCGAGATTTGGTTGTGCCTGGA |
| TTK reverse primer | CATCTGACACCAGAGGTTCCTTG |
| CDC20 forward primer | CGGAAGACCTGCCGTTACATTC |
| CDC20 reverse primer | CAGAGCTTGCACTCCACAGGTA |
| CENPC forward primer | GTGATGAGGCAGACTTGGCTAAG |
| CENPC reverse primer | TCCACTGGTCTGAGGCTTTAGG |
| DSN1 forward primer | TACCCAGTGCTTCCAGAAGGTG |
| DSN1 reverse primer | GTATGGCAGGTGGGTTCTGTAG |
| KIF18A forward primer | CTTGGACCAGTTCAGCCTATTCC |
| KIF18A reverse primer | GCACACTTTGAGATGGTGGAGAC |
| KIF23 forward primer | GTAGCAAGACCTGTAGACAAGGC |
| KIF23 reverse primer | TTCGCATGACGGCAAAGGTGGA |
| KIF14 forward primer | GCACTTTCGGAACAAGCAAACCA |

| | |
|---|---|
| KIF14 reverse primer | ATGTTGCTGGCAGCGGGACTAA |
| NCAPD2 forward primer | GTATGCTGCCTCTCATCTGGTC |
| NCAPD2 reverse primer | AGGCATCCACTAGCAGCAGAGA |
| KMT5A forward primer | GTGATTCCACCAATGCAGCCATC |
| KMT5A reverse primer | GCTCCTTCGGACAGGGTAGAAA |
| BUB1 forward primer | GCTCTGTCAGCAGACTTCCTTC |
| BUB1 reverse primer | CAGCAGATGTGAAGTCTCCTGG |
| CIAO1 forward primer | TTGGGTCTGGGAAGTTGATGA |
| CIAO1 reverse primer | CTCAAGGGTGGCACAGCATA |
| TRIP13 forward primer | ACTGTTGCACTTCACATTTTCCA |
| TRIP13 reverse primer | TCGAGGAGATGGGATTTGACT |
| CDC42 forward primer | TGACAGATTACGACCGCTGAGTT |
| CDC42 reverse primer | GGAGTCTTTGGACAGTGGTGAG |
| CAV1 forward primer | CCAAGGAGATCGACCTGGTCAA |
| CAV1 reverse primer | GCCGTCAAAACTGTGTGTCCCT |
| ANXA1 forward primer | GCGAAACAATGCACAGCGTCAAC |
| ANXA1 reverse primer | CAACCTCCTCAAGGTGACCTGT |
| RPL11 forward primer | AGAGTGGAGACAGACTGACGCG |
| RPL11 reverse primer | CGGATGCCAAAGGATCTGACAG |
| RPL26 forward primer | GGC TAA TGG CAC AAC TGT CCA C |
| RPL26 reverse primer | GGC GAG ATT TGG CTT TCC GTT C |
| RPL32 forward primer | ACA AAG CAC ATG CTG CCC AGT G |
| RPL32 reverse primer | TTC CAC GAT GGC TTT GCG GTT C |
| APC13 forward primer | GCT GCC TTA TGA GGA TGT CGC A |
| APC13 reverse primer | GGA ACA TTC TCA TGG AGG TAC TG |
| RPS27A forward primer | GCA GAG ACT GAT CTT TGC TGG C |
| RPS27A reverse primer | CTT GGG AGT GGT GTA AGA CTT CT |
| UBC forward primer | ACG GGA CTT GGG TGA CTC TA |
| UBC reverse primer | ATC GCC GAG AAG GGA CTA CT |
| ACTB forward primer | CCAACCGCGAGAAGATGA |
| ACTB reverse primer | CCAGAGGCGTACAGGGATAG |
| HPRT forward primer | TGACCTTGATTTATTTTGCATACC |
| HPRT reverse primer | CGAGCAAGACGTTCAGTCCT |
| Luciferase forward primer | CCAGGTATCAGGCAAGGATATG |
| Luciferase reverse primer | GTTCGTCTTCGTCCCAGTAAG |

**Table S10.** Overexpression vectors. Related to Figures 1-3.

| Vector | Source | Catalog number |
|---|---|---|
| pcDNA3 RLUC POLIRES FLUC | Addgene | 45642 |
| pT7-V5-SBP-C1-Hs4EBP1_Z | Addgene | 148150 |

| | | |
|---|---|---|
| eIF2a 1 | Addgene | 21807 |
| eIF2a 2 | Addgene | 21808 |
| pBrain-GFP-TACC3KDP-shTACC3 | Addgene | 59356 |
| pBrain-GFP-TACC3KDP(S558A)-shTACC3 | Addgene | 59357 |
| pRP[Exp]-CMV>{RPS27A 5'UTR}:Rluc | VectorBuilder | VB250812-1589pmd |
| pRP[Exp]-CMV>{HPRT 5'UTR}:Rluc | VectorBuilder | VB250813-1584ksk |
